# Invertebrate Community Associated with the Asexual Generation of *Bassettia pallida* Ashmead (Hymenoptera: Cynipidae)

**DOI:** 10.1101/2020.01.17.909648

**Authors:** Kelly L. Weinersmith, Andrew A. Forbes, Anna K.G. Ward, Pedro F. P. Brandão-Dias, Y. Miles Zhang, Scott P. Egan

## Abstract

Cynipid gall wasps play an important role in structuring oak invertebrate communities. Wasps in the Cynipini tribe typically lay their eggs in oaks (*Quercus* L.), and induce the formation of a “gall”, which is a tumor-like growth of plant material that surrounds the developing wasp. As the wasp develops, the cynipid and its gall are attacked by a diverse community of natural enemies, including parasitoids, hyperparasitoids, and inquilines. Determining what structures these species-rich natural enemy communities across cynipid gall wasp species is a major question in gall wasp biology. Additionally, gall wasps are ecosystem engineers, as the abandoned gall is used by other invertebrates. The gall-associated insect communities residing on live oaks (*Quercus geminata* Small and *Q. virginiana* Mill.) are emerging as a model system for answering ecological and evolutionary questions ranging from community ecology to the evolution of new species. Documenting the invertebrates associated with cynipids in this system will expand our understanding of the mechanisms influencing eco-evolutionary processes, record underexplored axes of biodiversity, and facilitate future work. Here, we present the community of natural enemies and other associates of the asexual generation of the crypt gall wasp, *Bassettia pallida* Ashmead. We compare the composition of this community to communities recently documented from two other cynipid gall wasps specializing on live oaks along the U.S. Gulf coast, *Disholcaspis quercusvirens* Ashmead and *Belonocnema treatae* Mayr. *B. pallida* and their crypts support a diverse arthropod community, including over 25 parasitoids, inquilines, and other associated invertebrates spanning 5 orders and 16 families.

---

Shelter-building insects (including gall formers, leaf rollers, leaf miners, and other insects that generate three-dimensional structures on their host plants) are ecosystem engineers, and are often associated with increases in arthropod richness and abundance on their host plants (reviewed in Cornelissen et al. 2016). While residing in their shelters, these insects are the target of parasitoids and predators, and are exploited by inquilines (i.e., organisms that typically do not make shelters themselves, but move into shelters with variable fitness implications for the shelter-maker) (Sanver and Hawkins 2000, Mendonça and Romanowski 2002, Hayward and Stone 2005, Bailey et al. 2009). The shelters often remain after the ecosystem engineer has abandoned it, and are subsequently settled by other arthropods (Cornelissen et al. 2016, Harvey et al. 2016, Wetzel et al. 2016).

Some of nature’s most complex shelters are created by cynipid gall wasps (Stone et al. 2002, Stone and Schönrogge 2003). Members of the Cynipini tribe lay their eggs in *Quercus* oaks (or sometimes other trees in the family Fagaceae), and induce the plant to produce a structure called a “gall”. The gall is lined with nutritious tissue, and will support the wasp while it feeds and develops (Rohfritsch 1992). Galls vary in appearance and structure, including those that exhibit exterior defenses that are sticky, hairy, or protruding spikes, or contain internal air sacs (Stone and Cook 1998, Csóka et al. 2005, Bailey et al. 2009), and those that are more cryptic (Melika and Abrahamson 2007). The incredible structural diversity among galls is thought to be a defense against the speciose community of parasitoids and inquilines that are often associated with galls (Abe et al. 2007, Askew et al. 2013), and which can have dramatic impacts on cynipid population fitness (Price et al. 1987, Sanver and Hawkins 2000, Stone and Schönrogge 2003, Csóka et al. 2005, Bailey et al. 2009). An estimated ∼1,400 described species of cynipid gall wasps produce morphologically diverse galls in their respective host plants (Ronquist et al. 2015, Pénzes et al. 2018), making this system ideal for asking questions about the importance of factors such as host relatedness, phenology, and natural enemy defense strategies on the structure of natural enemy communities (Cornell 1985, Stone et al. 2002, Stone and Schönrogge 2003, Price et al. 2004, Csóka et al. 2005, Hayward and Stone 2005).

Many of the studies on the structuring of the communities of natural enemies attacking cynipid gall wasps have been done in the Palaearctic (e.g., Schönrogge et al. 1996, Hayward and Stone 2005, Bailey et al. 2009, Nicholls et al. 2010, Stone et al. 2012, Bunnefeld et al. 2018). In North America, groundwork is being laid to conduct similar studies on the communities associated with the cynipid gall wasps that infect the “live oaks” (subsection Virentes) - a monophyletic group of seven North American semi-evergreen oak species within the genus *Quercus* – where much of the focus has centered on two partially overlapping sister species along the U.S. Gulf coast, *Quercus virginiana* and *Quercus geminata* (Cavender-Bares and Pahlich 2009, Cavender-Bares et al. 2015, Hipp et al. 2018). These live oaks are home to at least six (Egan et al. 2013) and potentially twelve (Egan, unpublished data) cynipid gall wasp species, which are in turn attacked by a community of parasitoids and inquilines (Bird et al. 2013, Forbes et al. 2016). The cynipid gall wasps and their communities of natural enemies associated with the live oak lineage are emerging as powerful systems for answering a broad set of ecological and evolutionary questions on local adaptation (Egan and Ott 2007), natural selection (Egan et al. 2011), speciation (Egan, Hood, and Ott 2012, Egan, Hood, Feder, et al. 2012, Egan et al. 2013, Zhang et al. 2017, 2019, Hood et al. 2019), developmental plasticity (Hood and Ott 2010), and novel species interactions (Egan et al. 2017, Weinersmith et al. 2017, Ward et al. 2019). However, the extent to which these species-rich natural enemy communities on gall wasp host species overlap is currently unclear, and even documentation of the species present in these communities is far from complete. Because of these limitations, the factors determining the overlap between the natural enemy communities attacking cynipid gall wasps species residing on live oaks remains unexplored.

After the host and/or its various natural enemies have exited the gall, the gall itself often remains. Abandoned oak galls are settled by a variety of arthropods, including ants, spiders, mites, and beetles (e.g., Cooper and Rieske 2010, Wetzel et al. 2016, Giannetti et al. 2019). While a recent review found that insect-made shelters are associated with increases in the abundance and diversity of arthropods that use the shelter once it is abandoned (Cornelissen et al. 2016), increases in density and diversity are not the rule. For example, the abandoned galls of the California gall wasp, *Andricus quercuscalifornicus* (Bassett), are associated with a *reduction* in herbivore invertebrate density and diversity, presumably because the galls are colonized by predatory spiders which attack herbivore invertebrates (Wetzel et al. 2016). Additional work is needed to better understand of the importance of abandoned galls on the local arthropod community.

Here we describe the community of natural enemies of the asexual generation of *Bassettia pallida* Ashmead (the crypt gall wasp), and discuss the overlap between the natural enemies of *B. pallida* and the previously described natural enemy communites of two gall wasps (*Belonocnema treatae* Mayr and *Disholcaspis quercusvirens* Ashmead) specializing on the same two live oak hosts, *Q. virginiana* and *Q. geminata*. The asexual generation of *B. treatae* creates leaf galls that contain one chamber, and was recently found to be associated with 24 invertebrate species (Forbes et al. 2016). The asexual generation of *D. quercusvirens* (which produces “bullet galls” on stems) was associated with nine species of parasitoids and inquilines (Bird et al. 2013).

Additionally, to stimulate future studies examining how *B. pallida* influences the diversity and abundance of the invertebrate community residing on live oaks, we report observations of associates of *B. pallida* that are likely benign and facultative, and use the crypt once it has been abandoned.

## Study System

The asexual generation of wasps in the North American genus *Bassettia* Ashmead (Hymenoptera, Cynipidae, Cynipini) produce stem galls in twigs, in which compartments where the wasps develop run parallel to the bark (Melika and Abrahamson 2007). The sexual generations of this genus – when known – make their galls in leaves, where they produce swellings that are visible on both sides of the leaf (Melika and Abrahamson 2007). The crypt gall wasp, *Bassettia pallida* (Hymenoptera: Cynipidae) (Fig. 1), infects both sand live oaks (*Q. geminata*) and southern live oaks (*Q. virginiana*) in the southeastern United States (Melika and Abrahamson 2007, Egan et al. 2013). The stem galls produced by the asexual generation of *B. pallida* are called “crypts”, and the sexual generation galls of this species have not been definitively identified. The community of natural enemies attacking *B. pallida* has not been described previously.

**Fig. 1.**
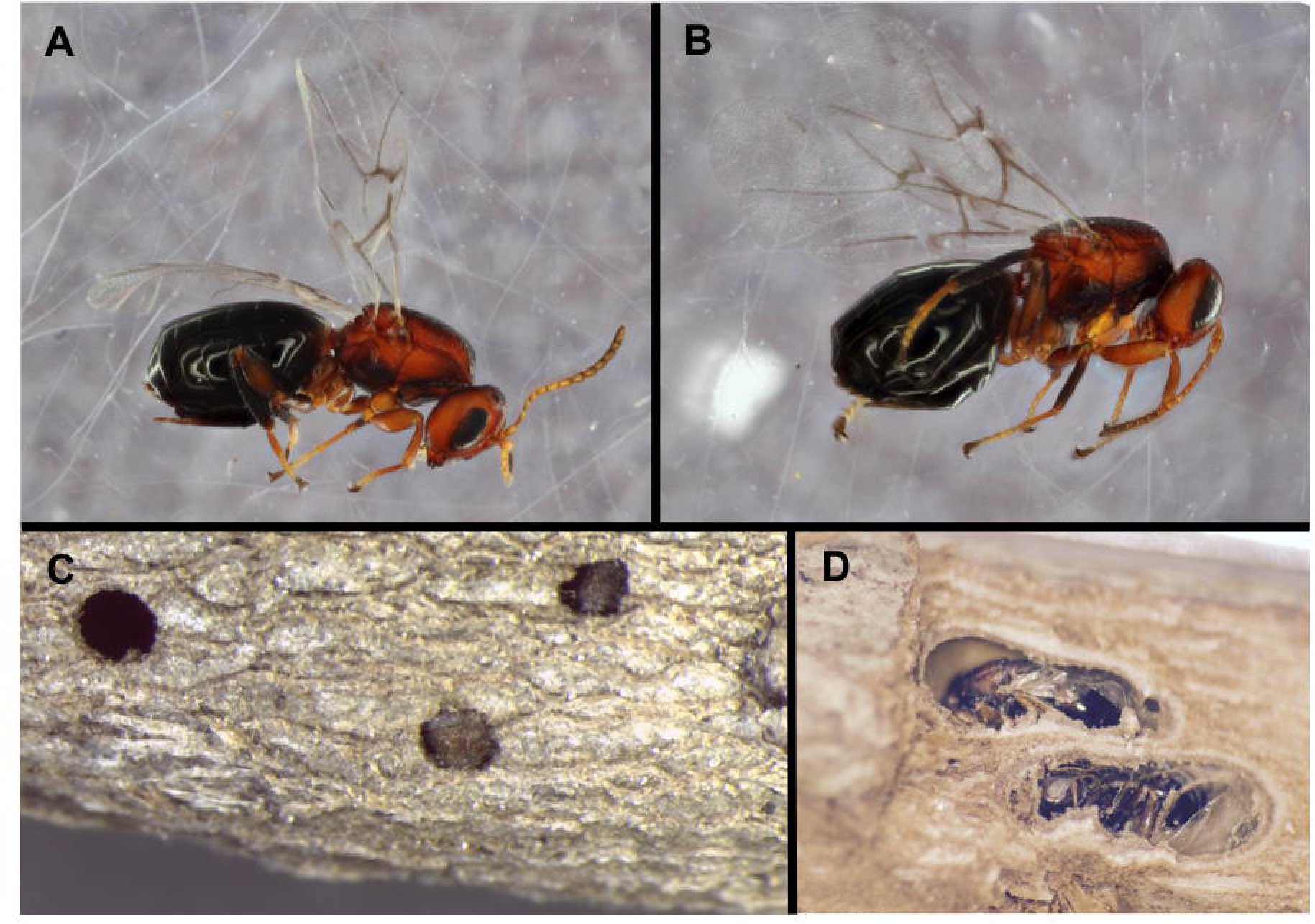
The asexual generation of *Bassettia pallida*, and their stem galls. (**A**) Female *B. pallida.* (**B**) Male *B. pallida*. (**C**) A *Quercus geminata* stem infected by *B. pallida*, showing the emergence hole from a crypt concealed with in the stem, and showing two *B. pallida* whose heads are plugging an incomplete emergence hole following manipulation by the parasitoid *Euderus set*. (**D**) Two crypt galls containing subadult *B. pallida*, revealed by removing the bark and some woody tissue using a razor blade. Photos A, B, and D originally appeared in Weinersmith et al. 2017, Proc Roy Soc B., and is available under a CC by 4.0 License.

## Materials and Methods

### Collections and Characterization of Natural Enemy Community

The stem galls made by the asexual generation of *Bassettia* are concealed, and are typically identified by finding emergence holes made by *Bassettia* which emerged previously (Melika and Abrahamson 2007). We collected *Q. geminata* stems with evidence of *B. pallida* emergence holes from four locations in Florida: Inlet Beach (Lat/Long: 30.273663, −86.001911), Lake Lizzie (28.227718, −81.179641), Topsail Hill Preserve State Park (30.3675327, −86.2752784), and Camp Helen State Park (30.270194, −85.991833). Collections made at Florida State Parks were made under Scientific Research Collecting Permit #04301840 from the Florida Department of Environmental Protection. We also collected *B. pallida*-infected *Q. virginiana* stems from two locations in Texas: Humble (29.998392, −95.184455) and Rice University’s Campus in Houston (29.717030, −95.401279). Collections occurred between August and March in 2015, 2016, 2018, and 2019. Tables 1 and 2 summarize collection years, locations sampled, host plant, and the number of invertebrates that emerged from each collection.

**Table 1:** Hymenopteran associates (including putative parasitoids, hyperparasitoids, and inquilines) of the asexual generation of *Bassettia pallida*. The table presents the number of specimens reared from live oak stems infected by *B. pallida* at various collection sites in Florida and Texas from 2015 – 2019. Collection site abbreviations: IB = Inlet Beach, LL = Lake Lizzie, TH = Topsail Hill Preserve State Park, CH = Camp Helen State Park, Rice U. = Rice University in Houston.

| Collection State, Host Tree<br>Collection Site<br>Collection Year (20XX) |  | Florida, *Quercus geminata* |  |  |  |  | Texas, *Quercus virginiana* |  |  |  |  |
| --- | --- | --- | --- | --- | --- | --- | --- | --- | --- | --- | --- |
|  |  | IB |  | LL | TH | CH | Humble |  |  | Rice U. |  |
|  |  | 15 | 19 | 15 | 18 | 19 | 15 | 16 | 19 | 16 | 17 |
| **Family** | **Species or subfamily** |  |  |  |  |  |  |  |  |  |  |
| Cynipidae | *Bassettia pallida* (galler) | 4 | 11 | 4 |  | 590 |  |  | 5 |  |  |
| Encyrtidae | *Encyrtidae* sp. |  |  |  |  | 1 |  |  |  |  |  |
| Eulophidae | *Euderus set* | 154 |  |  | 5 | 19 | 6 |  |  |  |  |
|  | *Galeopsomyia* sp. | 3 |  |  |  | 146 |  |  |  |  |  |
|  | *Tetrastichinae* sp. |  |  |  |  |  |  |  |  |  | 3 |
| Eupelmidae | *Brasema* sp. | 2 |  |  | 1 |  |  |  |  |  |  |
| Eurytomidae | *Eurytoma* sp. | 5 |  |  | 1 |  |  |  |  | 1 |  |
|  | *Sycophila* (4 morphotypes) | 13 |  |  |  |  |  | 1 |  | 2 | 7 |
| Ormyridae | *Ormyrus* nr. *thymus* | 5 |  |  |  | 1 |  |  |  |  |  |
|  | *Ormyrus* nr. *labotus* | 2 |  |  |  |  |  |  |  |  |  |
| Pteromalidae | *Acaenacis* sp. | 8 |  |  | 1 | 43 |  |  |  | 1 |  |
| Cynipidae | *Ceroptres* sp. | 1 |  |  |  |  |  |  |  |  |  |
|  | *Synergus walshii* | 1 |  |  |  |  |  |  |  |  |  |
| Formicidae | *Brachymyrmex obscurior* |  |  |  |  |  |  |  |  | 4 |  |
|  | *Crematogaster ashmeadi* |  |  | 8 |  |  |  |  |  |  |  |
| Braconidae | *Allorhogas* sp. |  |  |  |  | 8 |  |  |  |  |  |
| Platygastridae | *Telenomus* sp. | 1 |  |  |  |  |  |  |  |  |  |
|  | *Calotelea* sp. |  |  |  | 1 |  |  |  |  |  |  |
|  | *Synopeas* sp. | 3 |  |  |  |  |  |  |  |  |  |

**Table 2:** Associates of the asexual generation of *Bassettia pallida*. The table presents the number of specimens reared from live oak stems infected by *B. pallida* at various location sites in Florida (FL) and Texas (TX) from 2015 – 2019. Host tree abbreviations: *Qg* = *Quercus geminata*, *Qv* = *Quercus virginiana.* Collection site abbreviations: TH = Topsail Hill Preserve State Park, CH = Camp Helen State Park, Rice U. = Rice University in Houston.

| Collection State, Host Tree |  | FL, Qg |  |  |  |  | TX, Qv |  |
| --- | --- | --- | --- | --- | --- | --- | --- | --- |
| Collection Site |  | Inlet Beach |  |  | TH | CH | Rice U. |  |
| Collection Year (20XX) |  | 15 | 18 | 19 | 18 | 19 | 16 | 17 |
| **Order** |  |  |  |  |  |  |  |  |
| **Family** | **Subfamily/Species** |  |  |  |  |  |  |  |
| Coleoptera |  |  |  |  |  |  |  |  |
| Bostrichoidea | Ptinidae |  |  |  |  | 1 |  |  |
| Diptera |  |  |  |  |  |  |  |  |
| Cecidomyiidae | Cecidomyiinae sp 1 | 5 |  | 1 |  |  |  |  |
|  | Cecidomyiinae sp 2 |  |  |  |  |  |  | 1 |
| Psocoptera |  |  |  |  |  |  |  |  |
| Peripsocidae | *Peripsocus madidus* |  |  | 1 | 1 |  |  |  |
| Psocidae | Unknown |  |  |  |  | 2 |  |  |
| Lachesillidae | Unknown | 1 |  |  |  |  |  | 1 |
| Various | UnIDed Psocoptera | 3 | 2 | 7 | 13 | 2 | 2 | 1 |
| Thysanoptera |  |  |  |  |  |  |  |  |
| Phlaeothripidae | Unknown | 1 |  |  |  |  |  |  |

Stems collected in the field were placed in plastic bags and transported to either Rice University (Houston, Texas) or Charlottesville, Virginia. Leaves and non-target galls were removed from the stems, and stems were placed in clear plastic cups. The cups were covered with a coffee filter, which was secured in place with a rubber band. Cups were then placed outside, where they experienced natural light:dark cycles and ambient temperatures and humidity. The stems were misted with tap water periodically to mimic local precipitation. Abiotic differences between outdoor rearing conditions in Virginia and Texas may have influenced emergence success, but this is unlikely. After emergences ceased, haphazard dissections of stems suggested that most of the associates from the samples sent to Virginia had indeed emerged, and that no particular natural enemy species remained in the crypts. Cups were checked for emergences five days a week. Emerged insects were placed in 95% ethanol, and stored at room temperature or −20□ until further analysis.

Most emergent insects were Hymenoptera, which we identified using keys by Mason (1993), Gibson et al. (1997), Weld (1952), Gillette (1896), and Wahl (2019). For a subset of the associates, we extracted DNA using the DNeasy Blood and Tissue Kit (Qiagen) in accordance to the manufacturer’s protocol with the adition of a pestle crushing step prior to incubation. The mitochondrial cytochrome oxidase I (COI) region was amplified using the KAPA Taq ReadyMix (Sigma Aldrich) and the primers LEP F 5’ TAAACTTCTGGATGTCCAAAAAATCA 3’ and LEP R 5’ ATTCAACCAATACATAAAGATATTGG 3’ (Smith et al. 2008). Due to primer incompatibility, for the *Eurytoma* sample we used the following primers: COI_PF2 5’ ACC WGT AAT RAT AGG DGG DTT TGG DAA 3’ and COI_2437d 5’ CGT ART CAT CTA AAW AYT TTA ATW CCW G 3’ (Kaartinen et al. 2010). Thermocycling programs included 35 cycles with 48°C as the annealing temperature. We cleaned the resulting PCR products using the QIAquick PCR purification KIT (Qiagen) or an EXO1 (exonuclease 1) and SAP (shrimp alkaline phosphotase) method (15 min at 37°C minutes and then 80°C for an additional 15 min) prior to Sanger sequencing on an ABI 3730 (Applied Biosystems, Foster City, CA) in the University of Iowa’s Roy J. Carver Center for Genomics.

## Results

### Host Collection

*B. pallida* emerged from crypts in five of our collections, and emerged from both *Q. geminata* in Florida and *Q. virginiana* in Texas (Table 1). The greatest number of *B. pallida* emerged from crypts collected at Camp Helen State Park in Florida. Two *B. pallida* sequences (MN935926, MN935927; Table S1) were obtained from this location. The sequences were 98.98% identical, and multiple Cynipidae were ∼90-94% similar in GenBank. Most of the observed *B. pallida* emergences occurred in March, which is consistent with previous collections in Florida (Melika and Abrahamson 2007), and expectations from their natural history. Eight *B. pallida* emerged from late October through mid-December from collections made in the fall (August through October) and are likely responding to galled branches being removed from the tree.

Three specimens of an unidentified cynipid were found in the 2018-2019 collection. This cynipid appears to make crypt-like galls on stems and keys to the genus *Callirhytis* Foerster using (Zimmerman 2018), but could not be identified or matched to any currently described species. Upon further inspection, this ‘new’ species emerges from a solitary crypt gall with little to no external and visible swelling found at branching points within new stems, which is distinct from the cluster of crypt galls that generate a subtle swelling of the lateral parts of new branches induced by *B. pallida* (Brandão et al., MS in prep). We cannot rule out the possibility that some of the natural enemies and associates we describe below emerged from this galler. However, only 3 of the 590 (0.5%) cynipids that emerged from this collection were the non-target host species, suggesting that the vast majority of the natural enemies we collected were likely associated with *B. pallida*.

Associates from five orders and 16 families were reared from *Q. geminata* and *Q. virginiana* stems infected by the asexual generation of *B. pallida.* We present the Hymenopteran associates (Table 1) first, as they were the most abundant and diverse order present in our samples.

### Hymenoptera

We collected one specimen (Table 1) that keys to the family Encyrtidae (Chalcidoidea) using Grissell and Schauff (1997). The sequence from this specimen (MN935918, Table S1) was 97.7% similar to an unclassified Hymenopteran in GenBank. Encyrtid wasps can be parasitoids and hyperparasitoids, and many known host associations are with scale insects or mealybugs (Noyes 1988, Noyes and Woolley 1994). This wasp may be an associate of *B. pallida* galls, and not a direct parasitoid of the galler.

*Euderus set* Egan, Weinersmith, & Forbes (Chalcidoidea: Eulophidae; Fig 2A) emerged from *Q. virginiana* and *Q. geminata* at four sites. *E. set* is a recently described parasitoid of *B. pallida* (Egan et al. 2017), and manipulates its host into excavating an emergence hole from the crypt and then dying while plugging the hole with its head capsule (Fig. 1C, 2A) (Weinersmith et al. 2017). This behavior facilitates *E. set’s* escape from the crypt following completion of development (Weinersmith et al. 2017). Six other cynipid gall wasp hosts of *E. set* have recently been identified, all residing on different oak species than *B. pallida*, and all of which appear to be manipulated to facilitate parasitoid emergence (Ward et al. 2019).

**Fig. 2.**
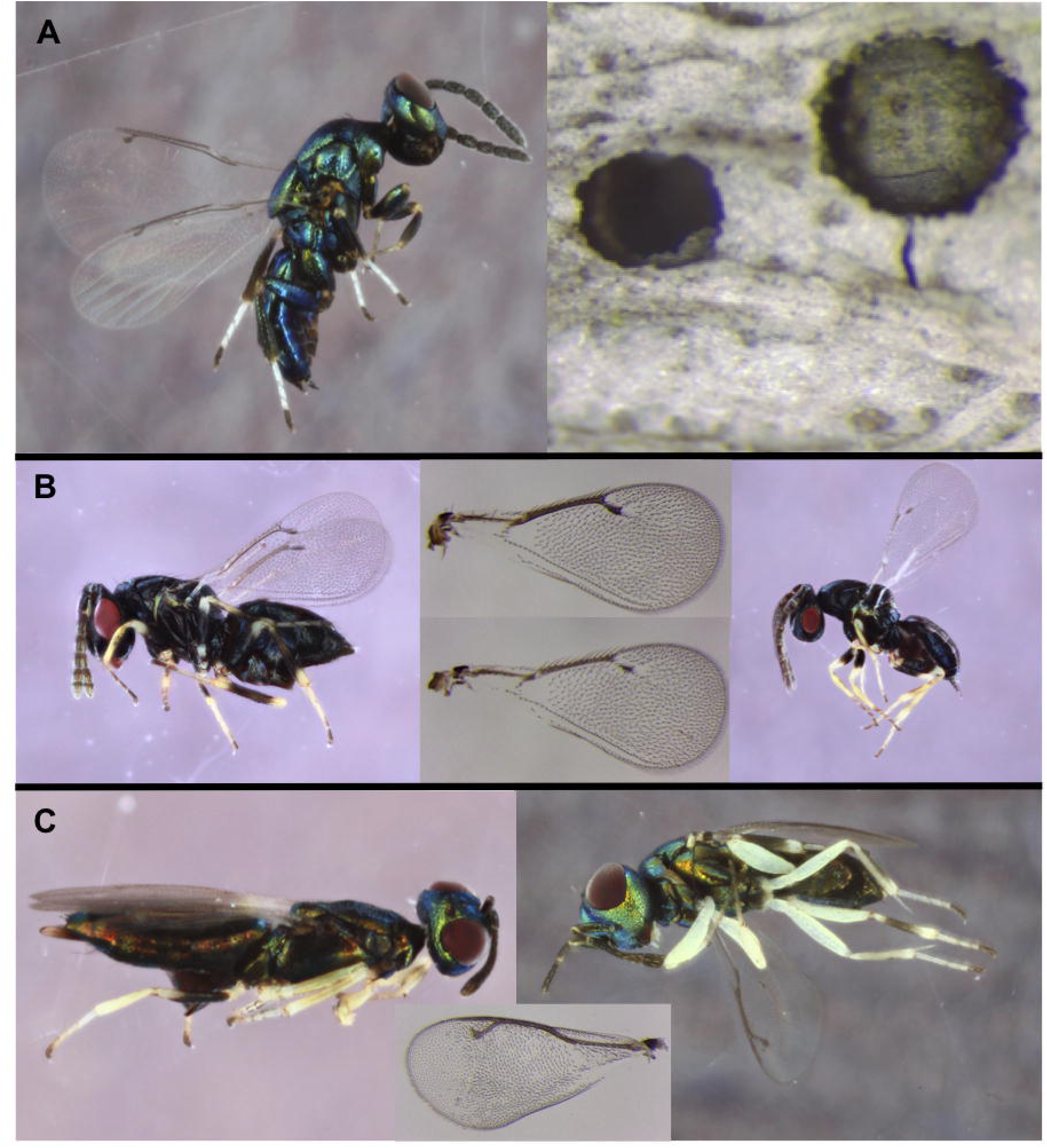
Natural enemies reared from *B. pallida* galls. (**A**) *Euderus set,* and examples of a *B. pallida* head capsule plugging an emergence hole (right) and a head-plugged emergence hole from which *E. set* has emerged (left). *E. set* photo originally appeared in Weinersmith et al. 2017, Proc Roy Soc B., and is available under a CC by 4.0 License. Photos of *B. pallida* head capsules by Mattheau Comerford. (**B**) *Galeopsomyia* species. Female on the left, with wing inset in top center. Male on the right, with wing inset in bottom center. (**C**) Unidentified *Brasema* species. Female on the left, male on the right, with an inset of a male’s wing in the center.

Two species of Tetrastichinae (Chalcidoidea: Eulophidae) emerged in our collections. The first species keyed to the genus *Galeopsomyia* Girault (Fig. 2B) using Schauff et al. (1997). This species emerged from three of our collections (Table 1), with the majority emerging mid-March through mid-April from the stems collected from *Q. geminata* at Camp Helen State Park (FL). We acquired a COI sequence (MN935919; Table S1), which was ∼84% similar to unclassified Eulophidae in GenBank. The second Tetrastichinae species was collected from *Q. virginiana* in Texas, and the sequence collected from this species (MN935910; Table S1) was only 79.9% similar to the *Galeopsomyia* species sequences. The tetrastichine wasp was 96.1% identical to an early release sequence from an Eulophidae in BOLD. In GenBank, this sequence was ∼86% identical to *Eulophidae* specimens in the subfamily Tetrastichinae.

Forbes et al. (2016) observed *Galeopsomyia nigrocyanea* (Ashmead) emerging from *B. treatae* on *Q. virginiana* in Texas. Based on sequence data, we suspect that the *Galeopsomyia* emerging from *B. pallida* are not *G. nigrocyanea*. The two *G. nigrocyanea* sequences deposited in GenBank from Forbes et al. (2016) are 84% similar to our Camp Helen specimen (from *Q. geminata* in Florida), and our Tetrastichinae specimen from Rice University (from *Q. virginiana* in Texas) shares only ∼83% similarity. Wasps in the genus *Galeopsomyia* are parasitoids of cynipid gall wasps (e.g., *Belonocnema treatae* (Forbes et al. 2016)), gall-forming dipterans (e.g., Cecidomyiidae (Stiling et al. 1992)), and are hyperparasitoids of other wasps, including some genera represented in our samples (e.g., *Eurytoma* Illiger (Herting 1977)). The *Galeopsomyia* that emerged from our samples could be a parasitoid, hyperparasitoid, or both (i.e., a facultative hyperparasitoid).

*Brasema* Cameron (Chalcidoidea: Eupelmidae: Eupelminae; Fig. 2C) were reared in our collections (Table 1), and were identified according to Gibson (1997). We obtained one sequence (MN935905; Table S1), which was 97.7% similar to “*Brasema* sp. GG5” (GenBank Accession HQ930308.1), which was collected by freehand sampling from a mesic hammock at Kissimmee Prairie Preserve State Park in Florida (BOLD Barcode Index Number BOLD:AAN7976). Based on sequence identity, the two *Brasema* species collected by Forbes et al. (2016) may be different species than that emerging from *B. pallida*. The sequences for *Brasema* sp. *1* and *Brasema* sp. *2* emerging from *B. treatae* (Forbes et al. 2016) are ∼89% and ∼91% similar to the sequence we collected from *Brasema* emerging from *B. pallida*. *Brasema* have a wide host range, with primary hosts including cynipid gall wasps, dipterans, and orthopterans, and a range of parasitoid hosts as well (including Pteromalids, Eurytomids, and Eulophids) (Noyes 2019). The exact relationship of *Brasema* to *B. pallida* is unknown.

*Eurytoma* (Chalcidoidea: Eurytomidae: Eurytominae; Fig. 3A) were reared from three collections (Table 1). Morphological ID of these specimens was done using DiGiulio (1997). We were unable to obtain sequence data for the one *Eurytoma* that emerged from two of the collections (one from *Q. virginiana* from Rice University in TX, and one from *Q. geminata* from Topsail Hill Preserve State Park in FL).

**Fig. 3.**
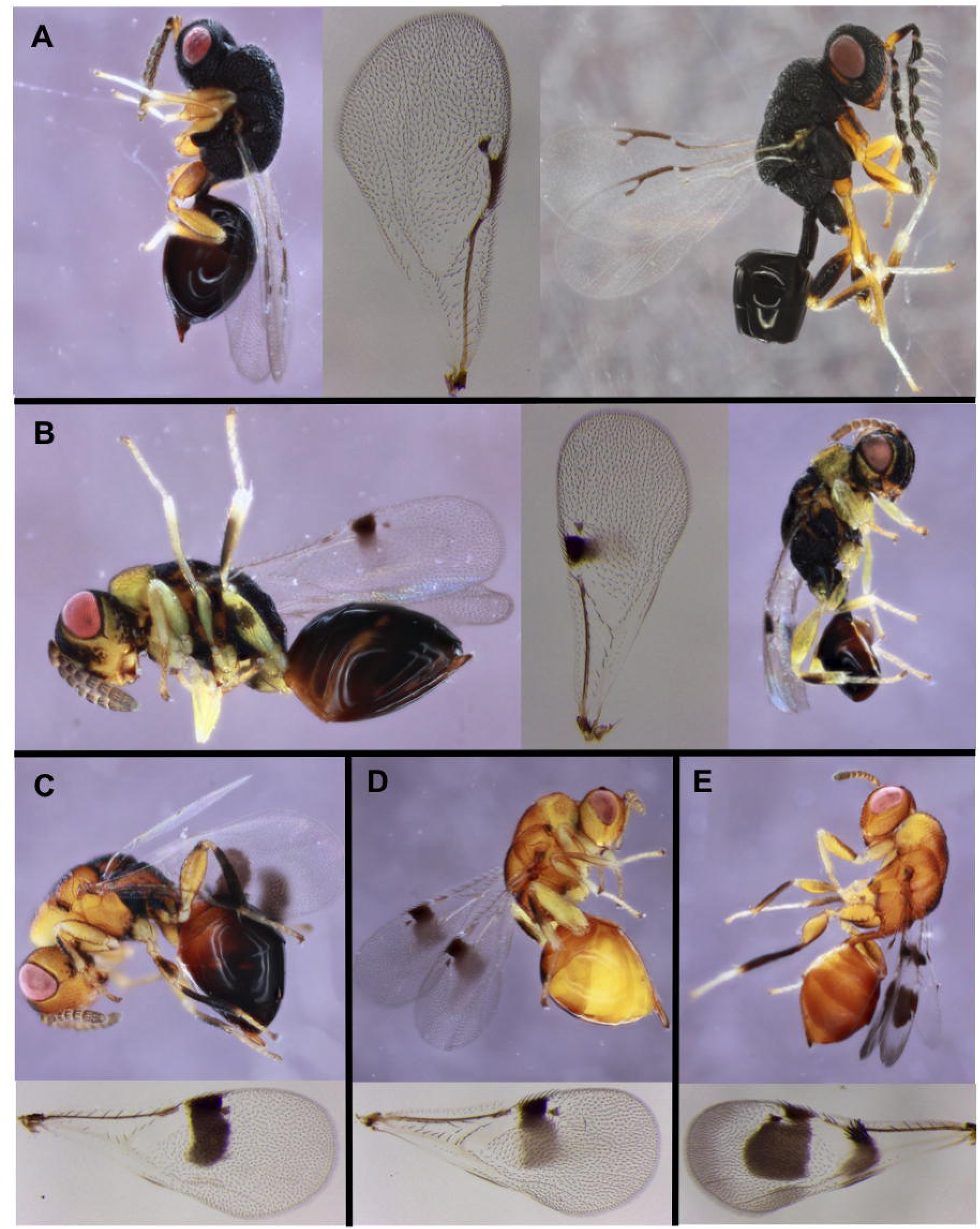
Eurytomidae natural enemies reared from galls made by the asexual generation of *B. pallida*. Males, when present, are on the right. Wings are from female specimens. (**A**) Unidentified *Eurytoma* species. (**B**) *Sycophila nr. foliatae*. (**C**) *Sycophila nr. dubia,* (**D**) *Sycophila nr. nubilistigma.* (**E**) *Sycophila nr. disholcaspidis*.

*Eurytoma* parasitoid species also emerge from galls of *B. treatae* and *D. quercusvirens*. Forbes et al. (2016) identified *Eurytoma furva* Bugbee and *Eurytoma bugbeei* Grissell, as well as one unidentified *Eurytoma* species emerging from *B. treatae.* Sequence comparisons between the *Eurytoma* specimen from *B. pallida* (MN935909; Table S1) and *E. furva* and *E. bugbeei* sequences collected from *B. treatae* (Forbes et al. 2016) are an ∼88% match, suggesting that the species collected from *B. pallida* is distinct. DNA could not be extracted from the unidentified *Eurytoma* species from *B. treatae*, so it is possible that this species emerges from both *B. pallida* and *B. treatae. Eurytoma hecale* Walker and an unidentified *Eurytoma* species were identified as parasitoids of *D. quercusvirens* by Bird et al. (2013). Our specimen is unlikely to be *E. hecale* based on morphology, and it is not possible to know if the unidentified *Eurytoma* species reported in Bird et al. (2013) is the same as that emerging from *B. pallida*.

Four morphospecies of *Sycophila* Walker (Chalcidoidea: Eurytomidae: Eurytominae, Fig. 3B-E) were reared from three collections (Table 1). The specimens were identified using Balduf (1923). The interpretation of the color variations in Balduf’s key is problematic, and the current species concepts are dubious until the taxonomic revision of the genus is conducted (Zhang et al., unpublished data). The first *Sycophila* morphospecies in our samples keys to *S. foliatae* (Ashmead) (Fig. 3B), which has previously been recorded from “live oak” in Jacksonville, FL, and is associated with a variety of oak gall parasitoids (Balduf 1923). The female specimens have varying degrees of black and yellow across the body and a small forewing infumation band, while males are mostly black with similar wing band patterns (Fig. 3B). These coloration characters also fit the description of *S. quinqueseptae* (Balduf), but this species is currently only known from California and is associated with *Plagiotrochus quinqueseptum* Ashmead (Balduf 1923). The second species identified was *S. nr. dubia* (Fig. 3C), although this species might be a synonym of *S. varians* based on morphology (large angular forewing infumation band, body color mix of yellow and black) and preliminary molecular studies (Zhang et al., unpublished data). *S. nr. nubilistigma* (Fig. 3D) is reared from *Q. virginiana*. They can be identified by their mostly yellow coloration with a dorsal black band on the mesoma and metasoma. The wing band is rectangular with a constriction near stigma vein (Fig. 3D). Finally, *S. nr. disholcaspidis* (Fig. 3E) was reared. The specimen is orange in coloration, and has the characteristic jug-shaped wing band and a secondary band near the parastigma similar to that of *S. disholcaspidis* Balduf which are parasitoids of *Disholcaspis cinerosa* (Bassett) in Texas. However, one key difference from *S. disholcaspidis* is the presence of multiple setae radiating from the secondary band (Fig. 3E), but more specimens are needed to better understand the species limits.

Three species of *Sycophila* (*S. texana* (Balduf), *S. varians* (Walsh), and *S. dorsalis* (Fitch)) were recently reared from the asexual generation of *D. quercusvirens* (Forbes et al. 2016), and one unidentified *Sycophila* species was reared from the asexual generation of *B. treatae* (Bird et al. 2013). While more work is needed to clarify the identities of these *Sycophila* species, it seems likely based on morphology that very little overlap occurs between the *Sycophila* species attacking *B. pallida, B. treatae,* and *D. quercusvirens*.

Two species of *Ormyrus* Westwood (Chalcidoidea: Ormyridae: Ormyrinae; Fig. 4A-B) emerged from crypts on *Q. geminata* from two of the Florida collections (Table 1), and were identified using the key in Hanson (1992). Two *Ormyrus nr. labotus* Walker (Fig. 4A) emerged from collections at Inlet Beach, FL. We obtained sequence data from one of these specimens (MN935904; Table S1), and the closest match in GenBank was to an unidentified Ormyridae. The sequence was also 91.5% to 93.6% identical to sequences from four *O. labotus* infecting *B. treatae* galls (Forbes et al. 2016). *O. labotus* is a generalist parasitoid, reported from more than 15 species of cynipid gall wasps (Noyes 2019). Additionally, six *O. nr. thymus* emerged (Fig. 4B, Table 1). A sequence obtained from one of these specimens (MN935907; TableS1) was an ∼89% match with an unidentified Ormyridae in BOLD. This sequence was only 86-87% similar to the *O. labotus* sequences associated with *B. treatae*. *Ormyus hegeli* (Girault) was also reared from both the asexual and sexual generation of *D. quercusvirens* (Bird et al. 2013), and based on morphology appears to be a different species from the two *Ormyrus* species reared from *B. pallida.* No known associates of *O. thymus* or *O. hegeli* are listed in Noyes (2019).

**Fig. 4.**
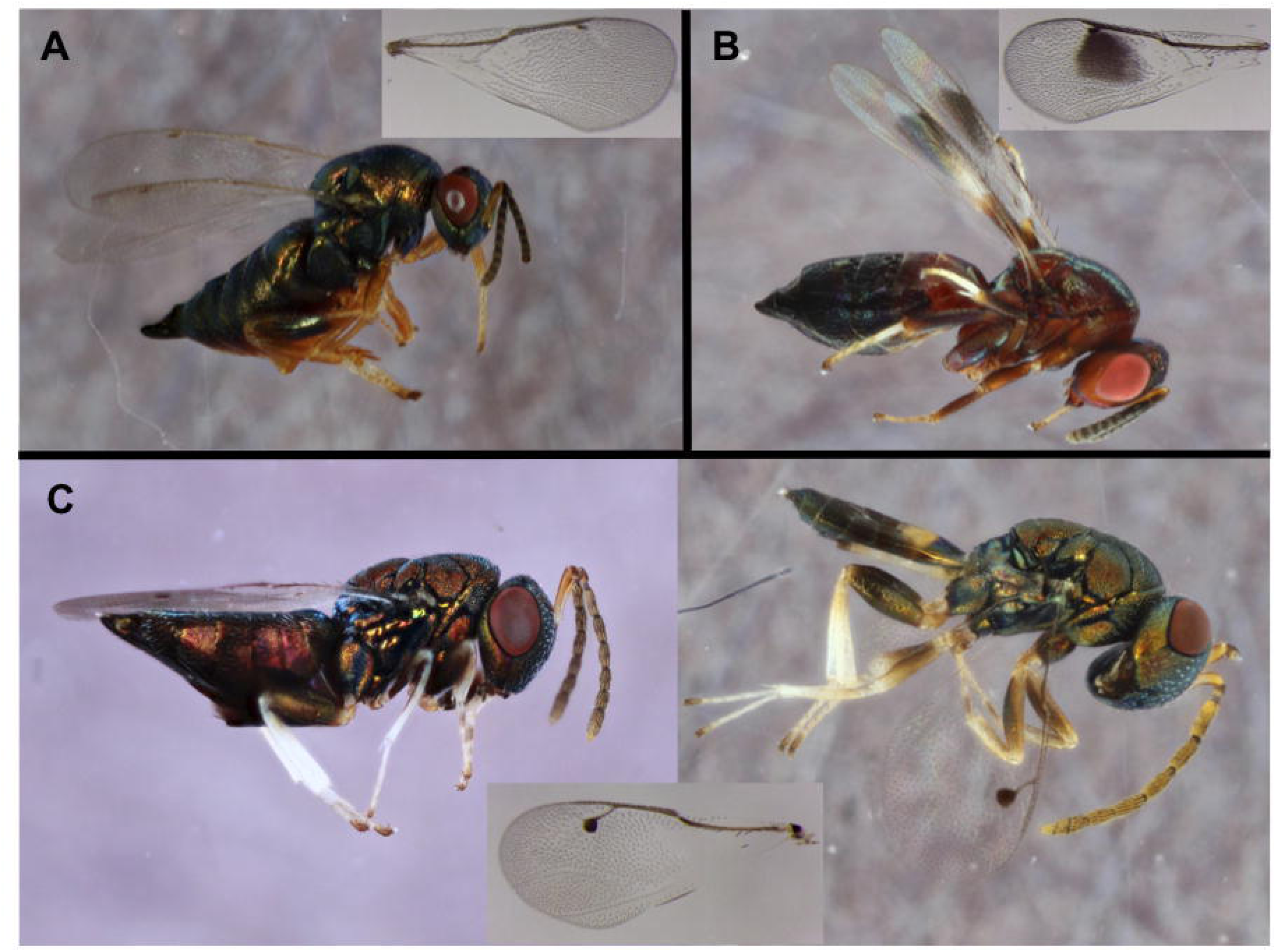
Natural enemies reared from *B. pallida* galls, with wing insets from female specimens. (**A**) *Ormyrus nr labotus*, (**B**) *Ormyrus nr thymus*, (**C**) Unidentified *Acaenacis* species, with female on left and male on right.

*Acaenacis* Girault (Chalcidoidea: Pteromalidae: Pteromalinae; Fig 4C) were reared from four sites, including stem galls from both *Q. virginiana* and *Q. geminata* (Table 1). These specimens keyed to the genus *Acaenacis* using Gibson et al. (1997). Three COI sequences (MN935908, MN935911, MN935912; TableS1) were 88.8% to 92.67% identical to each other, and the top hit for all three sequences in GenBank were to an unidentified Pteromalidae (85.5 to 88.3% identical).

Species in the genus *Acaenacis* attack oak-dwelling insects. *Acaenacis agrili* (Rohwer) is a parasitoid of the oak twig girdler (*Agrilus angelicus* Horn), which infects stems of *Quercus agrifolia* Nee in California (Rohwer 1919). Live oaks in the southeastern U.S. also harbor twig girdling beetles (Egan, S.P., personal observation). While associates of *Acaenacis taciti* (Girault) have not been identified (Noyes 2019), all other known hosts of *Acaenacis* are cynipid gall wasps. An undescribed *Acaenacis* infects *Andricus quercuslanigera* (Ashmead) on *Quercus rugosa* Nee in Mexico (Serrano-Muñoz et al. 2016). *Acaenacis lasus* (Walker) has been reared from leaf galls of *B. treatae* from *Quercus fusiformis* Small and *Q. virginiana* in Texas (Forbes et al. 2016), and *D. quercusvirens* on *Q. virginiana* in Florida (Bird et al. 2013). The four *A. lausus* sequences in GenBank from Forbes et al. (2016) were only 80-86% similar to the three *Acaenacis* sequences obtained from our collections. While *A. lasus* is infecting cynipid gall wasps on the same host plant as *B. pallida*, sequence data suggests that *A. lausus* is a different species than that emerging from *B. pallida*.

One *Ceroptres* sp. Hartig (Cynipoidea: Ceroptresini; Fig. 5A) was reared from a crypt collected at Inlet Beach, FL (Table 1), as was one *Synergus walshii* Gillette (Cynipoidea: Synergini; Fig. 5B). The *Ceroptres* specimen (MN935928; Table S1) was 90.4% similar to *Ceroptres* sp. *FSU 399* (Accession: DQ012636.1), which was reared from an *Andricus quercuscornigera* gall from Kentucky (USA) (Ronquist et al. 2015). The sequence from *S. walshii* (MN935929; TableS1) was 97.1% similar to *Synergus* sp. 1 from Forbes et al. (2016), which was one of three *Synergus* species associated with *B. treatae* in that study. Three *Synergus* are also associated with the asexual generation of *D. quercusvirens*, but it is unclear if the species in our study is the same as *Synergus* sp. 1 reported in Bird et al. (2013). Members of the genera *Synergus* and *Ceroptres* are inquilines of cynipid gall wasps, which are not able to initiate galls, but can maintain the production of nutritious tree tissue once inside a gall (Pénzes et al. 2012, Ronquist et al. 2015). *S. walshii* was reared from galls of several species of *Andricus* on various white oaks in IA, MO, and KY (A.K.G.W and A.A.F, unpublished data). Previous *S. walshii* collections were from *Andricus quercusflocci* galls on white oaks (*Quercus alba*) (Gillette 1896).

**Fig. 5.**
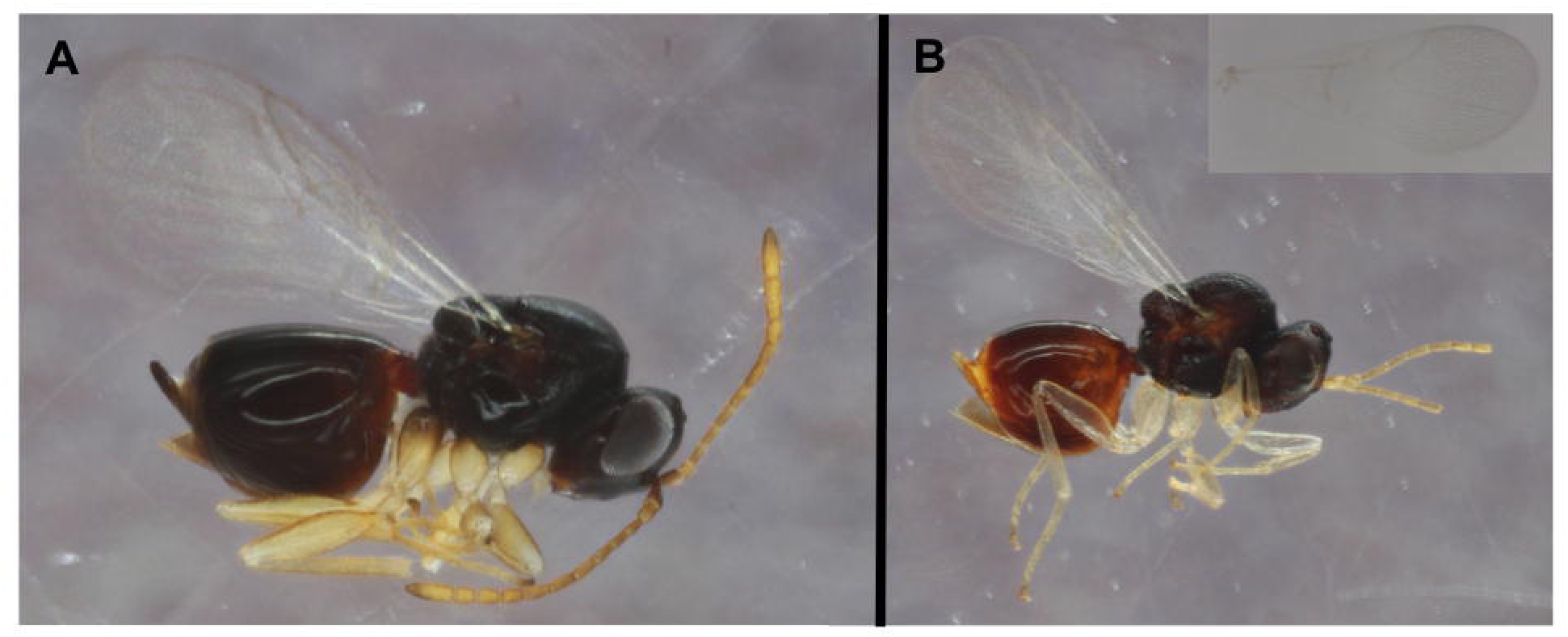
Inquilines reared from galls made by the the asexual generation of *B. pallida*. (**A**) Unidentified *Ceroptres.* (**B**) *Synergus walshii*, with inset showing detail of wing.

Ants (Formicidae) were observed twice during our samplings (Fig. 6, Table 1), and were identified to genus using Fisher and Cover (2007). Ants in the genus *Brachymyrmex* Mayr were observed and collected (N = 4 individuals; Fig. 6A) three months after *B. pallida*-infected *Q. virginiana* stems were brought into the lab. The sequence data from the *Brachymyrmex* (MN935915; TableS1) were 100% identical to a *Brachymyrmex obscurior* Forel sequence in GenBank. *Brachymyrmex obscurior* is likely an introduced species in North America (Deyrup et al. 2000). We also collected *Crematogaster ashmeadi* (N = 8 individuals; Fig. 6B), which were identified using Morgan and MacKay (2017). Some cynipid gall wasps induce their plant host to produce honeydew, which is consumed by ants who subsequently tend the gall (reviewed in Pierce 2019). While other cynipid gall wasps infecting oaks are known to secrete honeydew from their galls (e.g., *D. quercusvirens* on sand live oaks (Nicholls et al. 2017)), we did not observe honeydew on nor tending by ants of *B. pallida* crypts. Ants are also “secondary occupants” of galls, settling in the galls once cynipids have abandoned them (e.g., Giannetti et al. 2019). We suspect that *B. pallida-*abandoned crypts are occasionally settled by ants.

**Fig. 6.**
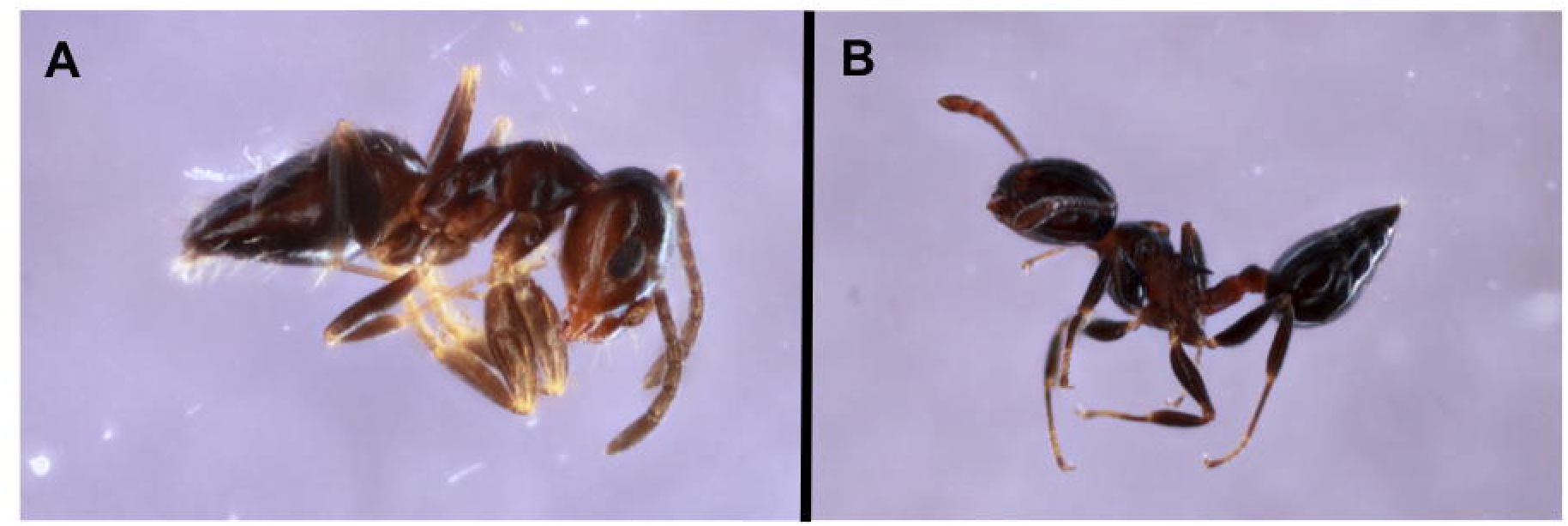
Ants (Formicidae) associated with *B. pallida* crypts. (**A**) *Brachymyrmex obscurior*. (**B**) *Crematogaster ashmeadi*.

The genus *Allorhogas* Gahan (Ichneumonoidea: Braconidae: Doryctinae) includes both gall-formers and parasitoids or inquilines (Zaldívar□Riverón et al. 2014). Eight *Allorhogas* (Table 1, Fig 7A) were reared from *Q. geminata*-infected stems from Camp Helen State Park (FL). A sequence was obtained from one of these specimens (MN935913; TableS1), and the sequence was 99.5% similar to a private *Allorhogas* sequence in BOLD. In GenBank, this sequence was 91.6% similar to the *Allorhogas* species reported from *B. treatae* in Forbes et al. (2016), and was 90.6 to 91.1% similar to an *Allorhogas* sp. 2 collected from South America (Zaldívar□Riverón et al. 2014). No *Allorhogas* were reported to infect *D. quercusvirens* in Bird et al. (2013). Whether this *Allorhogas* is a parasitoid, inquiline, or other associate of *B. pallida* is currently unknown, but the original description of the only *Allorhogas* species currently reported from the U.S. suggested that it might be a parasitoid of gall-associated lepidoptera burrowing through gall tissue (Gahan 1912).

**Fig. 7.**
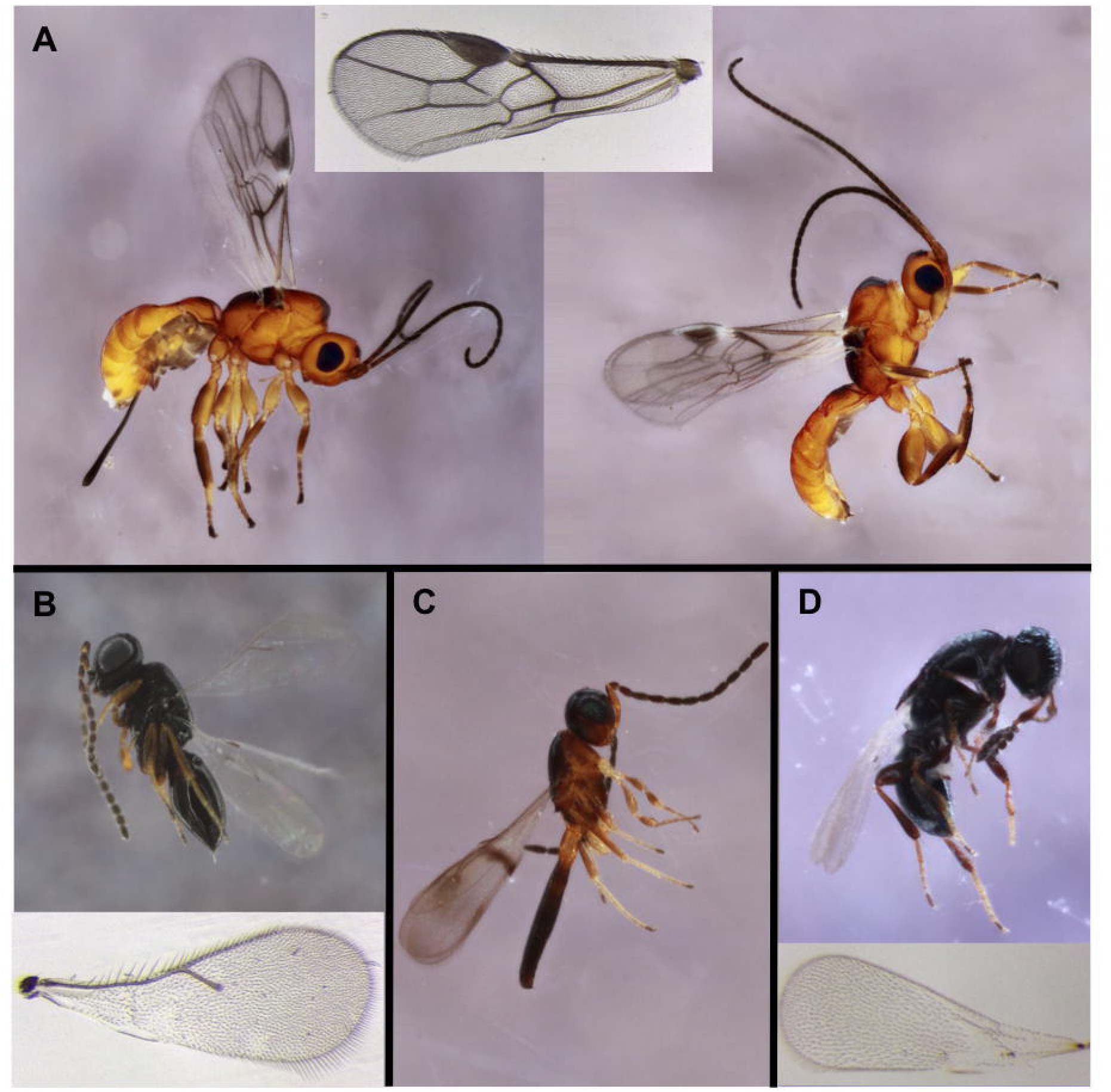
Natural enemies reared from galls made by the asexual generation of *B. pallida*, with wing insets from female specimens when available. (**A**) Unidentified *Allorhogas* species, with female on the left and male on the right, (**B**) Unidentified *Telenomus* species, (**C**) Unidentified *Calotelea* species, (**D**) Unidentified *Synopeas* species.

Five Platygastroidea specimens (Fig. 7B-D, Table 1) were identified using Mason (1993) and by Elijah Talamas (Florida Department of Agriculture and Consumer Services) based on specimen images. Sequence data were obtained from one specimen (MN935906; TableS1; Fig. 7B), which was ∼89% similar to an undescribed *Telenomus* Haliday species in BOLD and GenBank. The remaining four specimens belongs in the genus *Calotelea* Westwood and *Synopeas* Foerster (Scelionidae) (Fig. 7C and D, respectively). No Platygastridae or Scelionidae were observed emerging from the asexual generations of galls made by *B. treatae* (Forbes et al. 2016) or *D. quercusvirens* (Bird et al. 2013). Platygastroidea are typically egg parasitoids, and while specific Platygastroidea species often specialize, the range of hosts infected by parasitoids in this superfamily is broad (Murphy et al. 2007, Taekul et al. 2014). Both *Telenomus* and *Calotelea* may be egg parasitoids of *B. pallida* or other gall inhabitants, while *Synopeas* attacks the gall midge associate.

### Coleoptera

One beetle emerged from a *Q. geminata* stem in 2019 (Table 2, Fig. 8A), one month after the stem was brought into the lab. The sequence obtained from this beetle (MN935914; Table S1) was an ∼93% match to three published *Petalium bistriatum* (Ptinidae) sequences in BOLD. *Petalium* beetles are wood-boring (Ford 1973), and may not be directly associated with *B. pallida* crypts.

**Fig. 8.**
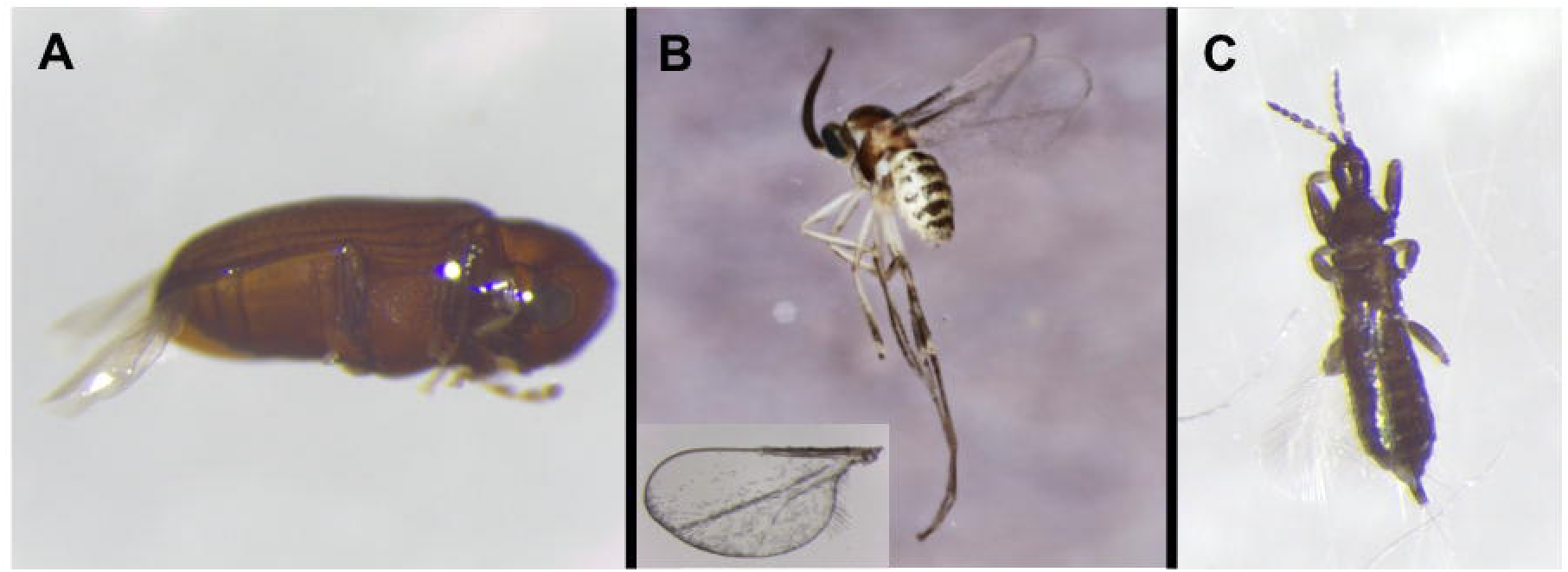
Associates of galls made by the asexual generation of *Bassettia pallida.* (**A**) Unidentified Ptinidae beetle (Coleoptera: Bostrichoidea), (**B**) Unidentified Cecidomyiidae (Diptera), (**C**) Unidentified Phlaeothripidae (Thysanoptera).

### Diptera

Two unidentified gall midge species in subfamily Cecidomyiinae (Sciaroidea: Cecidomyiidae) were reared in our collections (Table 2). The first species emerged from collections at Rice University’s campus on *Q. virginiana,* and the one sequence (MN935916; TableS1) obtained from this species is 99.9% similar to the unidentified gall midge associated with *B. treatae* (Forbes et al. 2016). The second species (MN935917; TableS1; Fig. 8B) was 91.9% similar to the first, emerged from Inlet Beach, FL on *Q. geminata,* and is ∼92% similar to *Asteromyia euthamiae* (subfamily Cecidomyiinae) sequences in GenBank. Gall midges are both gall formers and inquilines, including inquilines of cynipid gall wasps (Mamaev and Krivosheina 1992).

### Psocoptera

Thirty-six Psocopterans from at least three species emerged from the stems in our collections (Table 2, Fig 9). We sequenced six of these specimens, which we suspected represented two specimens for each of the three species. The rest of the Psocopterans we reared are reported as “Unidentified Psocoptera” in Table 2. Each pair of sequences from putative conspecifics were 99% similar to one another, and putative congener sequences were only ∼77-81% similar. Sequence data thus supports the presence of three Psocopteran species associated with *B. pallida* crypts. The sequences from the first species (MN935922 and MN935924; Table S1; Fig. 9A) were ∼98% similar to *Peripsocus madidus* sequences in GenBank. Two sequences from the second species (MN935923 and MN935925; Table S1; Fig. 9B) were ∼94% similar to an classified Psocodea in BOLD, and ∼84% similar to an unclassified Psocidae in GenBank. Two sequences (MN935920 and MN935921; TableS1) from the third species (Fig. 9C) were ∼97-99.7% similar to private Lachesillidae sequences in BOLD, and 83.1% similar to an unclassified Psocoptera in GenBank. These pscocopterans colonize abandoned crypts and inhabit them for long periods of time. They move in and out of the crypts, sometimes partially sealing the emergence holes with detritus.

**Fig. 9.**
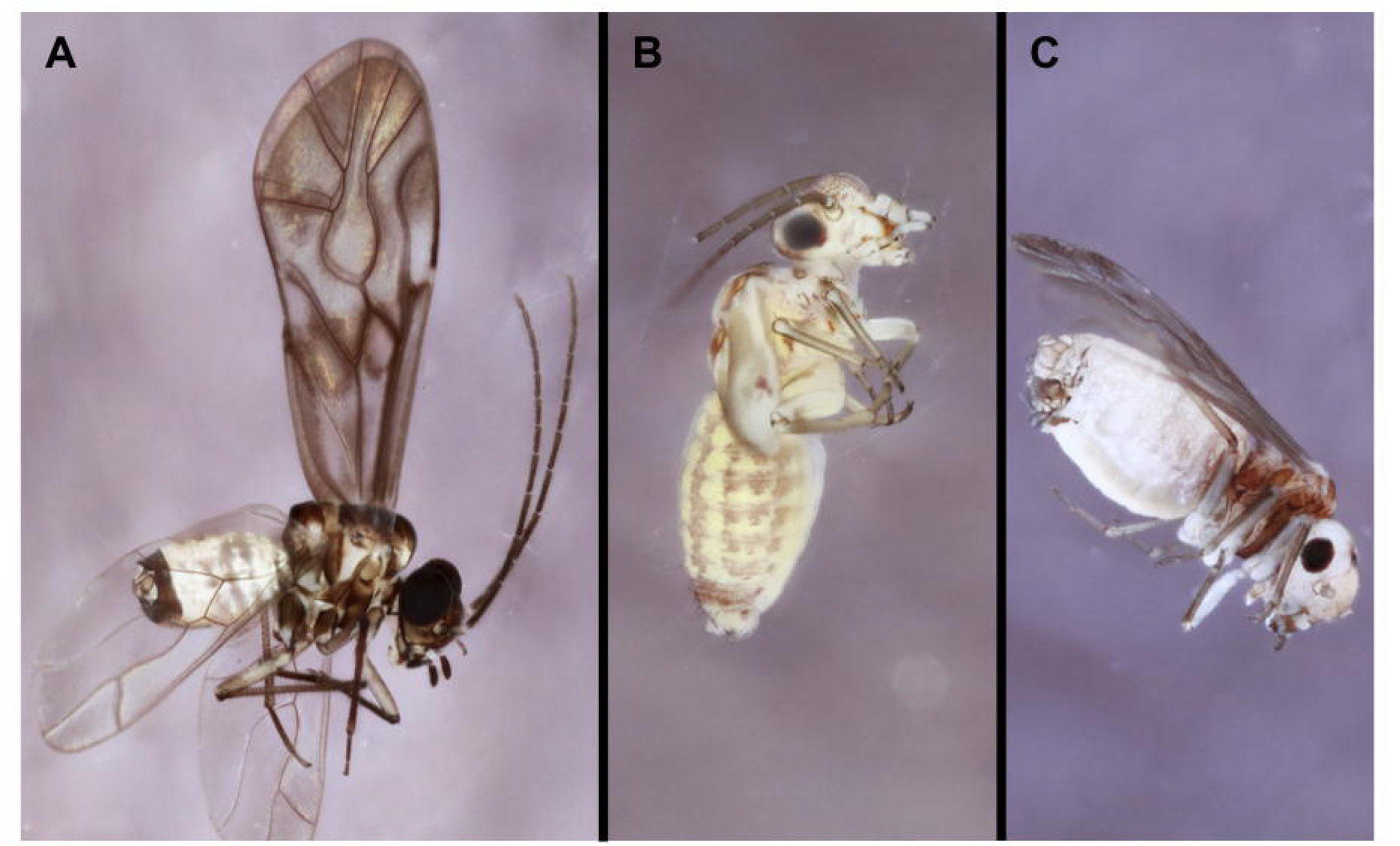
Psocopterans associated with galls made by the asexual generation of *Bassettia pallida*. (**A**) *Peripsocus madidus*. (**B**) Unidentified Psocidae. (**C**) Unidentified Lachesillidae.

### Thysanoptera

One thrips from the family Phlaeothripidae (Table 2, Fig. 8C) was associated with a stem collected in 2015 from *Q. geminata* at Inlet Beach, FL. We identified the specimen to family (Mound et al. 2009), but were unable to extract DNA.

## Discussion

A diverse and species-rich community of invertebrates is associated with the asexual generation of the gall wasp, *B. pallida*, and the crypts they create. These included Hymenoptera (21 species) (Table 1), Diptera (2), Coleoptera (1), Psocoptera (3), and Thysanoptera (1) (Table 2). The associates included parasitoids, inquilines, and secondary colonist that utilize the crypt after the emergence of *B. pallida* and/or its natural enemies.

### Putative Natural Enemies

Communities of natural enemies attacking cynipid gall wasps are structured by factors that include differences in gall structure and location on the host tree (e.g., leaf versus stem; Bailey et al. 2009). The natural enemy communities of three cynpid gall wasps residing on live oaks (*Q. geminata* and *Q. virginiana)* have been described recently (*D. quercusvirens*: Bird et al. 2013, *B. treatae:* Forbes et al. 2016, this study). The asexual generation of two of these wasps (*B. pallida* and *D. quercusvirens*) reside in similar locations (i.e., stems), while the asexual generation of *B. treatae* creates galls on leaves. The number of Hymenoptera that are likely parasitoids, hyperparasitoids, or inquilines was more similar between *B. pallida* (a stem-galler, ∼19 Hymenopteran natural enemies; this study) and *B. treatae* (a leaf-galler, ∼21 Hymenopteran natural enemies; (Forbes et al. 2016)), while the asexual generation of *D. quercusvirens* hosted only 9 natural enemies (Bird et al. 2013). While differences in sampling methods could explain this difference, it is also possible that *D. quercusvirens* had a low number of natural enemies due to its mutualism with ants. Some gall wasps attract ants by inducing the gall to produce honeydew, and these ants then defend the gall against inquilines and parasitoids (Washburn 1984, Abe 1992, Seibert 1993, Fernandes et al. 1999, Inouye and Agrawal 2004). Bird et al. (2013) noted the presence of ants on bullet galls created by *D. quercusvirens*, and these ants could have either excluded particular parasitoid species or reduced parasitoid success to low enough levels that these natural enemies were not observed during the collections.

While *B. pallida*, *D. quercusvirens*, and *B. treatae* all reside on live oaks, none of the natural enemies identified to species were common across all three cynipid gall wasp hosts. Additional natural enemies recorded for *D. quercusvirens* in Krombein et al. (1979) and for *B. treatae* in Peck (1963) overlap very little with natural enemies reported from the more recent studies. For *B. treatae*, only 1 of the 8 natural enemies reported in Peck (1963) was also reported by Forbes et al (2016), and, for *D. quercusvirens*, 7 natural enemies reported in Krombein et al. (1979) were not observed by Bird et al (2013). This suggests either that community composition is changing over time, that species were misidentified or the use of synonyms is confounding comparisons, or that no study has yet to sample these cynipid hosts with enough temporal and spatial coverage to capture the entire natural enemy community. However, there is some overlap between pairs of cynipid hosts. For example, *Acaenacis lausus* infects both *B. treatae* and *D. quercusvirens* (Bird et al. 2013, Forbes et al. 2016), and the inquiline *Synergus walshii* may be infecting both *B. treatae* and *B. pallida* (Forbes et al. 2016, and this study). In general though, it is difficult to draw strong conclusions about the degree of natural enemy overlap in these communities due to differences in sampling effort (including sampling done at different sites in different years). What is clear is that these galls support a diverse community of Hymenopteran natural enemies. As a rough estimation, if we assume that each of the 12 cynipid species on live oaks harbor about 15 host-specific natural enemies, that would yeild a community of 180 natural enemies in this system. Future work to quanitfy this diversity, and understand factors that influence the degree of overlap of these natural enemy communities across cynipid hosts is greatly needed.

One area of high interest is the association between *B. pallida* crypt galls and the recently described parasitoid *Euderus set*, which an example of a parasitoid species that can manipulate the behavior of its insect host (Weinersmith 2019). Specifically, E. set manipulates *B. pallida* into excavating an emergence hole from the crypt, which *B. pallida* then plugs with its head before being consumed by the parasitoid (Egan et al. 2017, Weinersmith et al. 2017). The parasitoid subsequently emerges from the host’s head capsule (Weinersmith et al. 2017). *E. set* infects and manipulates at least six additional cynipid gall wasp hosts infecting other oak species (Ward et al. 2019). The finding that *E. set* manipulates a broad range of cynipid gall wasps (Ward et al. 2019), suggests that the mechanism *E. set* uses to manipulate its host either does not require extreme specialization on host physiology or involves a mechanism common to many gall wasp residents. A more careful look at the gallers, inquilines, and parasitoids residing in live oak galls is warranted to determine if *E. set* is infecting and manipulating more hosts than just *B. pallida* in this system. Additionally, it is unclear why *E. set* is the only parasitoid that has been documented manipulating its hosts in this manner, while none of the other parasitoids or inquilines attacking cynipid gall wasps appear to do the same. Future work putting *E. set* in context with the other parasitoids infecting cynipid gall wasps should address questions about the selective pressure for manipulation, constraints on the evolution of this trait (including the costs paid by *E. set* as it manipulates its host), and the fitness benefits accrued through manipulation.

Finally, one common natural enemy that often attacks live oak galls was not observed in our study. Birds often break open the galls of *D. quercusvirens, D. cinerosa,* and *Callirhytis quercusbatatoides* (Ashmead) on live oaks to consume the wasps developing within (Weaver et al. in revision). During our collections we did not directly observe birds attacking *B. pallida* galls, nor did we see indirect evidence of predation on the stems. This suggests that the cryptic phenotype of *B. pallida* galls may to some extent protect the galler (and its natural enemy community) from bird predation.

### Other Associates

Associates reported in this study, which likely colonize *B. pallida* crypts after the galler, inquilines, and parasitoids have emerged, included ants, a beetle, a thrips, and barklice. While barklice were fairly common (36 were observed over the course of the study), the other associates were quite rare. Other reported colonizers of live oak galls include spiny millipedes (*Polyxenus* sp), spiders, mites, and lepidopterans (Wheeler and Longino 1988, Forbes et al. 2016). The lack of these associates in *B. pallida* galls could be explained by our sampling procedure underestimating the number of associates present, or because the small size of *B. pallida* crypts make them undesirable habitats for would-be colonizers. Ants, for example, seem to prefer to colonize larger galls (Almeida et al. 2014, Santos et al. 2017, Giannetti et al. 2019). The stem galls made by *B. pallida* tend to be smaller than the other cynipid galls on live oaks, and invertebrates may choose to colonize larger abandoned galls first.

We anticipated seeing lepidopterans in *B. pallida* galls, as they are associates of many galls on oaks (Brown and Mizell III 1993), including *B. treatae* leaf galls (Forbes et al. 2016) and *C. quercusbatatoides* stem galls (Egan, personal observation). The lack of lepidopterans has possible implications for our understanding of the biology of the parasitoid *Allorhogas,* as the original description of *Allorhogas* postulated that this genus may be parasitoids of lepidopterans (Gahan 1912). While it is possible that the timing of our collections missed lepidopterans associated with *B. pallida* crypts, the current data suggest that the *Allorhogas* observed in our system are not parasitoids of lepidopterans.

### Conclusions

While more detailed work may reveal some the associates we collected resided within the infected stem without actually associating with the crypt (e.g., this may be the case with the wood-boring Ptinid beetle), it is likely that future work will also reveal additional associates. In fact, our sampling may underestimate associate diversity in a number of ways. First, we have not sampled the unknown sexual generation of *B. pallida*, which remains to be discovered. This gall wasp generation will likely harbor some known species that attack both generations, as well as some that are unique to the sexual generation – as was found to be the case with the community attacking *B. treatae* on these same host plants (Forbes et al. 2016). Second, we identified infected live oak stems by looking for abandoned crypts, where these emergence holes were most likely formed in current and previous years. This highlights an important and general challenge to sampling cynipid associated communities, which is that time of sampling matters. We may have missed some parasitoids that emerge earlier - prior to the emergence of *B. pallida* – and/or missed some that attack later. Third, our sampling did not include the entire geographic range of *B. pallida*, which matches the distribution of its known host plant associations within the live oaks (*Q. virginiana* throughout the entire coastal southeastern United States from Virginina to Texas, *Q. geminata* restricted to xeric soils in Alabama, Mississippi, Florida, and Georgia, and *Q. fusiformis* in central and south Texas; see detailed host plant distributions in Cavender-Bares et al. 2015).

We found that the asexual generation of *B. pallida* is associated with a diverse arthropod community, including over 25 parasitoids, inquilines, and other invertebrates spanning five orders and 16 families. There was very little overlap between the natural enemy communities infecting two other live-oak infecting cynipid gall wasp species, suggesting a species-rich community of parasitoids and inquilines attacking cynipid gall wasps on live oaks. Descriptive studies like this are a necessary first step towards addressing broader ecological and evolutionary questions. In the future, we will use the community of cynipid gall wasps residing on live oaks, and the communities of natural enemies associated with these gallers, to address questions about habitat fragmentation and diversity (Maldonado-López et al. 2015), and the structuring of natural enemy communities (e.g., Bailey et al. 2009). Additionally, studies which quantify natural enemy communities and quantify the host range of parasitoids are critical for more accurate estimates of species richness (Forbes et al. 2018)

## Supporting information

Supplemental Table 1 GenBank Accession Numbers

## Acknowledgements

Linyi Zhang, Gaston Del Pino, Rebecca Izen, and Sean Liu assisted in regular cup checks, and we greatly appreciate their help. We thank Elijah Talamas for identifying the Platygastroidea specimens based on images. Thienthanh Trinh, Biplabendu Das, and Ian Will provided excellent company as well as assistance in the field during the 2018 collections. We also thank Clayton Iron Wolf at Camp Helen State Park for his assistance in finding live oaks at this site. KLW was funded by a Huxley Faculty Fellowship in Ecology and Evolutionary Biology from Rice University.

## References Cited

Abe, Y. 1992. The advantage of attending ants and gall aggregation for the gall wasp *Andricus symbioticus*(Hymenoptera: Cynipidae). Oecologia. 89: 166–167.

Abe, Y., G. Melika, and G. N. Stone. 2007. The diversity and phylogeography of cynipid gallwasps (Hymenoptera: Cynipidae) of the Oriental and eastern Palearctic regions, and their associated communities. Oriental Insects. 40: 169–212.

Almeida, M. F. B. de, L. R. dos Santos, and M. A. A. Carneiro. 2014. Senescent stem-galls in trees of *Eremanthus erythropappus* as a resource for arboreal ants. Revista Brasileira de Entomologia. 58: 265–272.

Askew, R. R., G. Melika, J. Pujade-Villar, K. Schönrogge, G. N. Stone, and J. L. Nieves-Aldrey. 2013. Catalogue of parasitoids and inquilines in cynipid oak galls in the West Palaearctic. Zootaxa. 3643: 1–133.

Bailey, R., K. Schönrogge, J. M. Cook, G. Melika, G. Csóka, C. Thuróczy, and G. N. Stone. 2009. Host niches and defensive extended phenotypes structure parasitoid wasp communities. PLoS Biology. 7: e1000179.

Balduf, W. V. 1923. Revision of the chalcid flies of the tribe Decatomini (Eurytomidae) in America north of Mexico. Proceedings of the US National Museum. 79: 38–41.

Bird, J. P., G. Melika, J. A. Nicholls, G. N. Stone, and E. A. Buss. 2013. Life history, natural enemies, and management of *Disholcaspis quercusvirens* (Hymenoptera: Cynipidae) on live oak trees. J Econ Entomol. 106: 1747–1756.

Brown, L. N., and R. F. Mizell III. 1993. The clearwing borers of Florida (Lepidoptera: Sesiidae). Tropical Lepidoptera. 4: 1–21.

Bunnefeld, L., J. Hearn, G. N. Stone, and K. Lohse. 2018. Whole-genome data reveal the complex history of a diverse ecological community. PNAS. 115: E6507–E6515.

Cavender-Bares, J., A. González-Rodríguez, D. A. R. Eaton, A. A. L. Hipp, A. Beulke, and P. S. Manos. 2015. Phylogeny and biogeography of the American live oaks (*Quercus* subsection Virentes): a genomic and population genetics approach. Molecular Ecology. 24: 3668–3687.

Cavender-Bares, J., and A. Pahlich. 2009. Molecular, morphological, and ecological niche differentiation of sympatric sister oak species, *Quercus virginiana* and *Q. geminata* (Fagaceae). American Journal of Botany. 96: 1690–1702.

Cooper, W. R., and L. K. Rieske. 2010. Gall structure affects ecological associations of *Dryocosmus kuriphilus* (Hymenoptera: Cynipidae). Environ Entomol. 39: 787–797.

Cornelissen, T., F. Cintra, and J. C. Santos. 2016. Shelter-building insects and their role as ecosystem engineers. Neotrop Entomol. 45: 1–12.

Cornell, H. V. 1985. Species assemblages of Cynipid gall wasps are not saturated. The American Naturalist. 126: 565–569.

Csóka, G., G. N. Stone, and G. Melika. 2005. Biology, ecology and evolution of gall-inducing Cynipidae, pp. 573–642. *In* Raman, A., Schaefer, C.W., Withers, T.W. (eds.), Biology, Ecology and Evolution of Gall-Inducing Arthropods.

Deyrup, M., L. Davis, and S. Cover. 2000. Exotic ants in Florida. Transactions of the American Entomological Society. 126: 293–326.

DiGiulio, J. A. 1997. Eurytomidae. *In* Gibson, G.A.P., Huber, J.T., Woolley, J.B. (eds.), Annotated Keys to the Genera of Nearctic Chalcidoidea (Hymenoptera). NRC Research Press.

Egan, S. P., G. R. Hood, G. DeVela, and J. R. Ott. 2013. Parallel patterns of morphological and behavioral variation among host-associated populations of two gall wasp species. PLOS ONE. 8: e54690.

Egan, S. P., G. R. Hood, J. L. Feder, and J. R. Ott. 2012. Divergent host-plant use promotes reproductive isolation among cynipid gall wasp populations. Biology Letters. 8: 605–608.

Egan, S. P., G. R. Hood, and J. R. Ott. 2011. Natural selection on gall size: Variable contributions of individual host plants to population-wide patterns. Evolution. 65: 3543– 3557.

Egan, S. P., G. R. Hood, and J. R. Ott. 2012. Testing the role of habitat isolation among ecologically divergent gall wasp populations. International Journal of Ecology. (https://www.hindawi.com/journals/ijecol/2012/809897/abs/).

Egan, S. P., and J. R. Ott. 2007. Host plant quality and local adaptation determine the distribution of a gall-forming herbivore. Ecology. 88: 2868–2879.

Egan, S. P., K. L. Weinersmith, S. Liu, R. D. Ridenbaugh, Y. M. Zhang, and A. A. Forbes. 2017. Description of a new species of *Euderus* Haliday from the southeastern United States (Hymenoptera, Chalcidoidea, Eulophidae): the crypt-keeper wasp. ZooKeys. 645: 37–49.

Fernandes, G. W., M. Fagundes, R. L. Woodman, and P. W. Price. 1999. Ant effects on three-trophic level interactions: plant, galls, and parasitoids. Ecological Entomology. 24: 411–415.

Fisher, B. L., and S. P. Cover. 2007. Ants of North America: a guide to the genera, First edition. ed. University of California Press, Berkeley.

Forbes, A. A., R. K. Bagley, M. A. Beer, A. C. Hippee, and H. A. Widmayer. 2018. Quantifying the unquantifiable: why Hymenoptera, not Coleoptera, is the most speciose animal order. BMC Ecology. 18: 21.

Forbes, A. A., M. C. Hall, J. Lund, G. R. Hood, R. Izen, S. P. Egan, and J. R. Ott. 2016. Parasitoids, hyperparasitoids, and inquilines associated with the sexual and asexual generations of the gall former, *Belonocnema treatae* (Hymenoptera: Cynipidae). Ann Entomol Soc Am. 109: 49–63.

Ford, E. J. 1973. A revision of the genus Petalium LeConte in the United States, Greater Antilles, and the Bahamas (Coleoptera:Anobiidae). U.S. Department of Agriculture.

Gahan, A. B. 1912. Descriptions of two new genera and six new species of parasitic Hymenoptera. Proceedings of the Entomological Society of Washington. 14: 2–8.

Giannetti, D., C. Castracani, F. A. Spotti, A. Mori, and D. A. Grasso. 2019. Gall-colonizing ants and their role as plant defenders: from ‘bad job’ to ‘useful service.’ Insects. 10: 392.

Gibson, G. A. P., J. T. Huber, and J. B. Woolley. 1997. Annotated keys to the genera of nearctic Chalcidoidea (Hymenoptera). NRC Research Press.

Gillette, C. P. 1896. A monograph of the genus *Synergus* Hartig. Transactions of the American Entomological Society (1890-). 23: 85–100.

Hanson, P. 1992. The Nearctic species of *Ormyrus* Westwood (Hymenoptera: Chalcidoidea: Ormyridae). Journal of Natural History. 26: 1333–1365.

Harvey, J. A., P. J. Ode, M. Malcicka, and R. Gols. 2016. Short-term seasonal habitat facilitation mediated by an insect herbivore. Basic and Applied Ecology. 17: 447–454.

Hayward, A., and G. N. Stone. 2005. Oak gall wasp communities: evolution and ecology. Basic and Applied Ecology, Special feature: Gall-Inducing Insects - Nature’s Most Sophisticated Herbivores. 6: 435–443.

Herting, B. 1977. Hymentoptera. A catalogue of parasites and predators of terrestrial arthropods. Section A. Host or prey/enemy. Commongwealth Agricultural Bureaux, Institute of Biological Control, Farnham Royal, England.

Hipp, A. L., P. S. Manos, A. González-Rodríguez, M. Hahn, M. Kaproth, J. D. McVay, S. V. Avalos, and J. Cavender-Bares. 2018. Sympatric parallel diversification of major oak clades in the Americas and the origins of Mexican species diversity. New Phytologist. 217: 439–452.

Hood, G. R., and J. R. Ott. 2010. Developmental plasticity and reduced susceptibility to natural enemies following host plant defoliation in a specialized herbivore. Oecologia. 162: 673– 683.

Hood, G. R., L. Zhang, E. G. Hu, J. R. Ott, and S. P. Egan. 2019. Cascading reproductive isolation: plant phenology drives temporal isolation among populations of a host-specific herbivore. Evolution. 73: 554–568.

Inouye, B. D., and A. A. Agrawal. 2004. Ant mutualists alter the composition and attack rate of the parasitoid community for the gall wasp *Disholcaspis eldoradensis* (Cynipidae). Ecological Entomology. 29: 692–696.

Kaartinen, R., G. N. Stone, J. Hearn, K. Lohse, and T. Roslin. 2010. Revealing secret liaisons: DNA barcoding changes our understanding of food webs. Ecological Entomology. 35: 623–638.

Krombein, K. V., P. D. Hurd, D. R. Smith, and B. D. Burks. 1979. Catalog of Hymenoptera in America north of Mexico, vol. 1: Symphyta and Aprocrita (Parasitica). Smithsonian Institution Press, Washington, D.C.

Maldonado-López, Y., P. Cuevas-Reyes, G. N. Stone, J. L. Nieves-Aldrey, and K. Oyama. 2015. Gall wasp community response to fragmentation of oak tree species: importance of fragment size and isolated trees. Ecosphere. 6: art31.

Mason, W. R. M. 1993. Key to superfamily of Hymenoptera, pp. 65–100. In Goulet, H., Huber, J.T. (eds.), Hymentoptera of the World: An Identification Guide to Families. Agriculture Canada, Ottawa, Ontario.

Melika, G., and W. G. Abrahamson. 2007. Review of the Nearctic gallwasp species of the genus *Bassettia* Ashmead, 1887, with description of new species (Hymenoptera: Cynipidae: Cynipini). Acta Zoologica Academiae Scientiarum Hungaricae. 53: 131–148.

Mendonça, M. de S., and H. P. Romanowski. 2002. Natural enemies of the gall-maker *Eugeniamyia dispar* (Diptera, Cecidomyiidae): predatory ants and parasitoids. Brazilian Journal of Biology. 62: 269–275.

Morgan, C., and W. Mackay. 2017. The North America acrobat ants of the hyperdiverse genus Crematogaster. LAP LAMBERT Academic Publishing.

Mound, L. A., D. Paris, and N. Fisher. 2009. Phlaeothripidae. World Thysanoptera. (http://anic.ento.csiro.au/thrips/identifying_thrips/Phlaeothripidae.htm).

Murphy, N. P., D. Carey, L. R. Castro, M. Dowton, and A. D. Austin. 2007. Phylogeny of the platygastroid wasps (Hymenoptera) based on sequences from the 18S rRNA, 28S rRNA and cytochrome oxidase I genes: implications for the evolution of the ovipositor system and host relationships. Biol J Linn Soc. 91: 653–669.

Nicholls, J. A., P. Fuentes-Utrilla, A. Hayward, G. Melika, G. Csóka, J.-L. Nieves-Aldrey, J. Pujade-Villar, M. Tavakoli, K. Schönrogge, and G. N. Stone. 2010. Community impacts of anthropogenic disturbance: natural enemies exploit multiple routes in pursuit of invading herbivore hosts. BMC Evolutionary Biology. 10: 322.

Nicholls, J. A., G. Melika, and G. N. Stone. 2017. Sweet tetra-trophic tnteractions: multiple evolution of nectar secretion, a defensive extended phenotype in Cynipid gall wasps. The American Naturalist. 189: 67–77.

Noyes, J. S. 1988. Encyrtidae (Insecta: Hymenoptera), Fauna of New Zealand. DSIR Science Information Publishing Centre, Wellington.

Noyes, J. S. 2019. Universal Chalcidoidea Database. (http://www.nhm.ac.uk/chalcidoids).

Noyes, J. S., and J. B. Woolley. 1994. North American encyrtid fauna (Hymenoptera: Encyrtidae): taxonomic changes and new taxa. Journal of Natural History. 28: 1327– 1401.

Peck, O. 1963. A catalogue of the nearctic Chalcidoidea (Insecta: Hymenoptera). Memoirs of the Entomological Society of Canada. 95: 1–1092.

Pénzes, Z., C.-T. Tang, P. Bihari, M. Bozsó, S. Schwéger, and G. Melika. 2012. Oak associated inquilines (Hymenoptera, Cynipidae, Synergini), TISCIA monograph series. Szeged.

Pénzes, Z., C.-T. Tang, G. N. Stone, J. A. Nicholls, S. Schwéger, M. Bozsó, and G. Melika. 2018. Current status of the oak gallwasp (Hymenoptera: Cynipidae: Cynipini) fauna of the Eastern Palaearctic and Oriental Regions. Zootaxa. 4433: 245–289.

Pierce, M. P. 2019. The ecological and evolutionary importance of nectar-secreting galls. Ecosphere. 10: e02670.

Price, P. W., W. G. Abrahamson, M. D. Hunter, and G. Melika. 2004. Using gall wasps on oaks to test broad ecological concepts. Conservation Biology. 18: 1405–1416.

Price, P. W., G. W. Fernandes, and G. L. Waring. 1987. Adaptive nature of insect galls. Environ Entomol. 16: 15–24.

Rohfritsch, O. 1992. Patterns in gall development, pp. 60–86. *In* Shorthouse, J.D., Rohfritsch, O. (eds.), Biology of Insect-Induced Galls. Oxford University Press, New York, N.Y.

Rohwer, S. A. 1919. Descriptions of three parasites of *Agrilus angelicus* (Hym.). Proceedings of the Entomological Society of Washington. 21: 4–8.

Ronquist, F., J.-L. Nieves-Aldrey, M. L. Buffington, Z. Liu, J. Liljeblad, and J. A. A. Nylander. 2015. Phylogeny, Evolution and Classification of Gall Wasps: The Plot Thickens. PLOS ONE. 10: e0123301.

Santos, L. R. dos, R. dos S. M. Feitosa, and M. A. A. Carneiro. 2017. The role of senescent stem-galls over arboreal ant communities structure in *Eremanthus erythropappus* (DC.) MacLeish (Asteraceae) trees. Sociobiology. 64: 7–13.

Sanver, D., and B. A. Hawkins. 2000. Galls as habitats: the inquiline communities of insect galls. Basic and Applied Ecology. 1: 3–11.

Schauff, M. E., J. LaSalle, and L. D. Coote. 1997. Eulophidae, pp. 327–429. *In* Gibson, G.A.P., Huber, J.T., Wooley, J.B. (eds.), Annotated Keys to the Genera of Neartic Chalcidoidae (Hymenoptera). NRC Research Press, Ottawa, Ontario.

Schönrogge, K., G. N. Stone, and M. J. Crawley. 1996. Alien herbivores and native parasitoids: rapid developments and structure of the parasitoid and inquiline complex in an invading gall wasp *Andricus quercuscalicis* (Hymenoptera: Cynipidae). Ecological Entomology. 21: 71–80.

Seibert, T. F. 1993. A nectar-secreting gall wasp and ant mutualism: selection and counter-selection shaping gall wasp phenology, fecundity and persistence. Ecological Entomology. 18: 247–253.

Serrano-Muñoz, M., G. A. Villegas-Guzmán, A. Callejas-Chavero, J. R. Lomeli-Flores, U. M. Barrera-Ruíz, J. Pujade, and M. Ferrer-Suay. 2016. Hymenopterans associated with *Andricus quercuslanigera* galls (Hymenoptera: Cynipidae, Chalcidoidea) from sierra de Guadalupe, State of México. Entomología Mexicana. 177–182.

Smith, M. A., J. J. Rodriguez, J. B. Whitfield, A. R. Deans, D. H. Janzen, W. Hallwachs, and P. D. N. Hebert. 2008. Extreme diversity of tropical parasitoid wasps exposed by iterative integration of natural history, DNA barcoding, morphology, and collections. Proceedings of the National Academy of Sciences. 105: 12359–12364.

Stiling, P., A. M. Rossi, D. R. Strong, and D. M. Johnson. 1992. Life history and parasites of *Asphondylia borrichiae* (Diptera: Cecidomyiidae), a gall maker on *Borrichia frutescens*. The Florida Entomologist. 75: 130–137.

Stone, G. N., and J. M. Cook. 1998. The structure of cynipid oak galls: patterns in the evolution of an extended phenotype. Proceedings of the Royal Society of London. Series B: Biological Sciences. 265: 979–988.

Stone, G. N., K. Lohse, J. A. Nicholls, P. Fuentes-Utrilla, F. Sinclair, K. Schönrogge, G. Csóka, G. Melika, J.-L. Nieves-Aldrey, J. Pujade-Villar, M. Tavakoli, R. R. Askew, and M. J. Hickerson. 2012. Reconstructing community assembly in time and space reveals enemy escape in a western palearctic insect community. Current Biology. 22: 532–537.

Stone, G. N., and K. Schönrogge. 2003. The adaptive significance of insect gall morphology. Trends in Ecology & Evolution. 18: 512–522.

Stone, G. N., K. Schönrogge, R. J. Atkinson, D. Bellido, and J. Pujade-Villar. 2002. The population biology of oak gall wasps (Hymenoptera: Cynipidae). Annual Review of Entomology. 47: 633–668.

Taekul, C., A. A. Valerio, A. D. Austin, H. Klompen, and N. F. Johnson. 2014. Molecular phylogeny of telenomine egg parasitoids (Hymenoptera: Platygastridae s.l.: Telenominae): evolution of host shifts and implications for classification. Systematic Entomology. 39: 24–35.

Wahl, D. B. 2019. Genera Ichneumonorum Nearcticae. (http://www.amentinst.org/GIN/).

Ward, A. K. G., O. S. Khodor, S. P. Egan, K. L. Weinersmith, and A. A. Forbes. 2019. A keeper of many crypts: a behaviour-manipulating parasite attacks a taxonomically diverse array of oak gall wasp species. Biology Letters. 15: 20190428.

Washburn, J. O. 1984. Mutualism between a cynipid gall wasp and ants. Ecology. 65: 654– 656.

Weaver, A. K., G. R. Hood, M. Foster, and S. P. Egan. in revision. The shape and magnitude of phenotypic selection at different spatial scales is driven by bottom up and top down effects.

Weinersmith, K. L. 2019. What’s gotten into you?: a review of recent research on parasitoid manipulation of host behavior. Current Opinion in Insect Science, Pests and resistance • Behavioural ecology. 33: 37–42.

Weinersmith, K. L., S. M. Liu, A. A. Forbes, and S. P. Egan. 2017. Tales from the crypt: a parasitoid manipulates the behaviour of its parasite host. Proc. R. Soc. B. 284: 20162365.

Weld, L. H. 1952. Cynipoidea (Hym.) 1905-1950, being a supplement to the Dalla Torre and Kieffer Monograph: The Cynipidae in Das Tierreich, Lieferung 24, 1910, and bringing the systematic literature of the world Up to date, including keys to families and subfamilies and lists of new generic, specific and variety names. Privately printed.

Wetzel, W. C., R. M. Screen, I. Li, J. McKenzie, K. A. Phillips, M. Cruz, W. Zhang, A. Greene, E. Lee, N. Singh, C. Tran, and L. H. Yang. 2016. Ecosystem engineering by a gall-forming wasp indirectly suppresses diversity and density of herbivores on oak trees. Ecology. 97: 427–438.

Wheeler, J., and J. T. Longino. 1988. Arthropods in live oak galls in Texas. Entomological News. 99: 25–29.

Zaldívar-Riverón, A., J. J. Martínez, S. A. Belokobylskij, C. Pedraza-Lara, S. R. Shaw, P. E. Hanson, and F. Varela-Hernández. 2014. Systematics and evolution of gall formation in the plant-associated genera of the wasp subfamily Doryctinae (Hymenoptera: Braconidae). Systematic Entomology. 39: 633–659.

Zhang, L., A. Driscoe, R. Izen, C. Toussaint, J. R. Ott, and S. P. Egan. 2017. Immigrant inviability promotes reproductive isolation among host-associated populations of the gall wasp *Belonocnema treatae*. Entomologia Experimentalis et Applicata. 162: 379–388.

Zhang, L., G. R. Hood, J. R. Ott, and S. P. Egan. 2019. Temporal isolation between sympatric host plants cascades across multiple trophic levels of host-associated insects. Biology Letters. 15: 20190572.

Zimmerman, J. R. 2018. A synopsis of oak gall wasps (Hymenoptera: Cynipidae) of the southwestern United States with a key and comments on each of the genera. Journal of the Kansas Entomological Society. 91: 58–70.

