## Supplemental Table 1 GenBank Accession Numbers for "Invertebrate Community Associated with the Asexual Generation of *Bassettia pallida* Ashmead (Hymenoptera: Cynipidae)"

**Supplemental Table 1 –** GenBank Accession Numbers for COI sequences collected for this study. Order, family, species or subfamily (when known), lab code, harvest date (the date when the crypt was collected from the field), location where collection occurred, and the host plant (*Quercus geminata* or *Quercus virginiana*) on which the crypt was found is included.

| **GenBank #** | **Order** | **Family** | **Species or Subfamily** | **Lab Code** | **Harvest Date** | **Location** | **Host plant** |
| --- | --- | --- | --- | --- | --- | --- | --- |
| MN935914 | Coleoptera | Ptinidae |  | 216 | 18-Mar-19 | Camp Helen State Park, FL | *Q. geminata* |
| MN935916 | Diptera | Cecidomyiidae | Cecidomyiinae | SL128 | 23-Jan-17 | Houston, TX (Rice University) | *Q. virginiana* |
| MN935917 | Diptera | Cecidomyiidae | Cecidomyiinae | 220 | 18-Mar-19 | Inlet Beach, Florida | *Q. geminata* |
| MN935913 | Hymenoptera | Braconidae | *Allorhogas sp.* | 214 | 18-Mar-19 | Camp Helen State Park, FL | *Q. geminata* |
| MN935926 | Hymenoptera | Cynipidae | *Bassettia pallida* | LZ6423_1 | 18-Mar-19 | Inlet Beach, Florida | *Q. geminata* |
| MN935927 | Hymenoptera | Cynipidae | *Bassettia pallida* | LZ6423_2 | 18-Mar-19 | Inlet Beach, Florida | *Q. geminata* |
| MN935928 | Hymenoptera | Cynipidae | *Ceroptres sp.* | SL110 | 15-Oct-15 | Inlet Beach, Florida | *Q. geminata* |
| MN935929 | Hymenoptera | Cynipidae | *Synergus walshii* | SL002 | 1-Aug-15 | Inlet Beach, Florida | *Q. geminata* |
| MN935918 | Hymenoptera | Encyrtidae |  | 211 | 18-Mar-19 | Camp Helen State Park, FL | *Q. geminata* |
| MN935910 | Hymenoptera | Eulophidae | Tetrastichinae | KW113 | 23-Jan-17 | Houston, TX (Rice University) | *Q. virginiana* |
| MN935919 | Hymenoptera | Eulophidae | *Galeopsomyia sp.* | 145 | 18-Mar-19 | Camp Helen State Park, FL | *Q. geminata* |
| MN935905 | Hymenoptera | Eupelmidae | *Brasema sp.* | KW002 | 1-Aug-15 | Inlet Beach, Florida | *Q. geminata* |
| MN935909 | Hymenoptera | Eurytomidae | *Eurytoma sp.* | SL056 | 1-Aug-15 | Inlet Beach, Florida | *Q. geminata* |
| MN935915 | Hymenoptera | Formicidae | *Brachymyrmex obscurior* | SL126 | 13-Nov-16 | Houston, TX (Rice University) | *Q. virginiana* |
| MN935904 | Hymenoptera | Ormyridae | *Ormyrus sp. nr. labotus* | SL001 | 1-Aug-15 | Inlet Beach, Florida | *Q. geminata* |
| MN935907 | Hymenoptera | Ormyridae | *Ormyrus sp. nr. thymus* | KW004 | 1-Aug-15 | Inlet Beach, Florida | *Q. geminata* |
| MN935906 | Hymenoptera | Platygastridae | *Telenomus* | SL017 | 1-Aug-15 | Inlet Beach, Florida | *Q. geminata* |
| **GenBank #** | **Order** | **Family** | **Species or Subfamily** | **Lab Code** | **Harvest Date** | **Location** | **Host plant** |
| MN935908 | Hymenoptera | Pteromalidae | *Acaenacis sp.* | KW003 | 1-Aug-15 | Inlet Beach, Florida | *Q. geminata* |
| MN935911 | Hymenoptera | Pteromalidae | *Acaenacis sp.* | 154 | 18-Mar-19 | Camp Helen State Park, FL | *Q. geminata* |
| MN935912 | Hymenoptera | Pteromalidae | *Acaenacis sp.* | 6 | 8-Sep-18 | Topsail Hill Preserve State Park, Florida | *Q. geminata* |
| MN935920 | Psocoptera | Lachesillidae |  | KW112 | 16-Jan-17 | Houston, TX (Rice University) | *Q. virginiana* |
| MN935921 | Psocoptera | Lachesillidae |  | KW099 | 15-Oct-15 | Inlet Beach, Florida | *Q. geminata* |
| MN935922 | Psocoptera | Peripsocidae | *Peripsocus madidus* | 225 | 18-Mar-19 | Inlet Beach, Florida | *Q. geminata* |
| MN935924 | Psocoptera | Peripsocidae | *Peripsocus madidus* | 21 | 8-Sep-18 | Topsail Hill Preserve State Park, Florida | *Q. geminata* |
| MN935923 | Psocoptera | Psocidae |  | 102 | 18-Mar-19 | Camp Helen State Park, FL | *Q. geminata* |
| MN935925 | Psocoptera | Psocidae |  | 118 | 18-Mar-19 | Camp Helen State Park, FL | *Q. geminata* |
